# CGRP Administration into the Cerebellum Evokes Migraine-like Behaviors Predominately in Female Mice

**DOI:** 10.1101/2022.01.24.477577

**Authors:** Mengya Wang, Thomas L. Duong, Brandon J. Rea, Jayme S. Waite, Michael W. Huebner, Harold C. Flinn, Andrew F. Russo, Levi P. Sowers

## Abstract

The neuropeptide calcitonin gene-related peptide (CGRP) is a major player in migraine pathophysiology. Previous preclinical studies demonstrated that intracerebroventricular administration of CGRP caused migraine-like behaviors in mice, but the sites of action in the brain remain unidentified. The cerebellum has the most CGRP binding sites in the central nervous system and is increasingly recognized as both a sensory and motor integration center. The objective of this study was to test whether the cerebellum, particularly the medial cerebellar nuclei (MN), might be a site of CGRP action. In this study, CGRP was directly injected into the right MN of C57BL/6J mice via a cannula. A battery of behavioral tests was done to assess migraine-like behaviors. CGRP caused light aversion measured as decreased time in the light zone even with dim light. The mice also spent more time resting in the dark zone, but not the light, along with decreased rearing and transitions between zones. These behaviors were similar for both sexes. In contrast, significant responses to CGRP were seen only with female mice in the open field assay, von Frey test, and automated squint assay, indicating anxiety, tactile hypersensitivity, and spontaneous pain, respectively. In male mice, the responses had the same trend as females but did not reach statistical significance. No detectable effect of CGRP on gait was observed in either sex. These results suggest that CGRP in the MN causes light aversion in males, while in females, light aversion is accompanied by increased anxiety, tactile hypersensitivity, and spontaneous pain. A caveat is that we cannot exclude contributions from other cerebellar regions in addition to the MN due to diffusion of the injected peptide. These results reveal the cerebellum as a new site of CGRP actions that may contribute to migraine pathophysiology and possibly its prevalence in females.

## 1 Introduction

Migraine is a neurological disease that affects about 15% of the population (1) and is the second leading cause of disability globally (2). It is characterized by moderate or severe headaches that are accompanied by sensory abnormalities, such as photophobia and allodynia (3). Prevalence in women is about twice as high as in men (1). Despite its high prevalence and large burden to society, the mechanism underlying migraine have yet to be fully elucidated. Over the last few decades, calcitonin gene-related peptide (CGRP) has moved to the forefront in migraine pathophysiology. CGRP levels are elevated in both the ictal and interictal phases in human studies (4-6) and infusion of CGRP induced migraine-like headaches in ∼66% of migraine patients (7-10). Most recently, CGRP-based drugs have been shown to effectively alleviate migraine symptoms in about 50% of patients (11, 12). However, despite the significant advancement of CGRP-based drugs as migraine therapeutics, there is uncertainty regarding the mechanisms by which CGRP induces migraine, especially as to where CGRP is acting.

The human studies measuring CGRP levels (4-6) and induction of migraine-like headaches by intravenous CGRP injections (7-10) suggest a peripheral site of action for CGRP in migraine. In addition, the antibodies targeted against CGRP or CGRP receptors have limited ability to cross the blood-brain barrier (13). However, previous animal studies demonstrated that peripheral (intraperitoneal, i.p.) (14) and central (intracerebroventricular, i.c.v.) (15) injection of CGRP induced similar light-aversive behaviors in wild-type mice. Both behaviors could be attenuated by triptan migraine drugs (14, 15). Moreover, transgenic mice overexpressing a CGRP receptor subunit in the nervous system displayed light aversion in response to dim light after i.c.v. CGRP injection(14, 16, 17), while bright light was required to induce light aversion in wild-type mice after i.c.v. CGRP injection (15). Those data suggest that increased sensitivity to CGRP in the nervous system can cause migraine-like light-aversive behavior in mice. Finally, it was found that CGRP injection into the posterior thalamic nuclei, an integration center for light and pain signals, was sufficient to induce light aversion in wild-type C57BL/6J mice, even in dim light (18). Together, these data suggest that CGRP can work in the central nervous system to induce migraine-like photophobic behavior in mice.

Similar to the thalamus, the cerebellum integrates multiple sensory signals and motor events (19, 20). While the cerebellum was originally recognized for its role in motor control (21), there is mounting evidence that it also plays important roles in perceptual (22), emotional (23), and cognitive functions (24-26). In particular, it is now appreciated that the cerebellum participates in sensory, emotional, cognitive aspects of pain, and motor control in response to pain (27). Three lines of evidence support the link between the cerebellum and migraine pathogenesis. First, changes in cerebellar activation (28-30), structure (31-35) (36-38) (39-43), and functional connectivity (44-49) are present in episodic, chronic, and familial hemiplegic migraine patients. When responding to trigeminal stimuli, cerebellar activity and functional connectivity with the thalamus and cortical areas were changed (34), suggesting the cerebellum is involved in processing sensory information from the trigeminal system. Strikingly, migraine patients exhibited decreased cerebellar activation in response to trigeminal nociceptive stimuli after treatment with erenumab, a CGRP receptor antibody (50). Second, migraine patients display cerebellar symptoms, e.g., dizziness, vertigo (51), body sway (52, 53), as well as increased body sway accompanied by increased light intensity (54). Third, the cerebellum communicates directly to migraine-related regions, such as the spinal trigeminal nucleus (55-57) and the thalamus (58) via direct neural circuits. These data hint to the importance of the cerebellum in migraine pathophysiology.

Curiously, the cerebellum has the highest binding density to CGRP receptor PET ligands in human and rhesus brains (59, 60). The canonical CGRP receptor subunits, receptor activity-modifying protein 1 (RAMP1) and calcitonin receptor-like receptor (CLR), are localized in the human, rhesus and rat cerebellar cortex (61-66) and in the medial cerebellar nuclei (MN, also known as fastigial nuclei in humans) of rats (65). CGRP is also distributed in the cerebellar cortex (61, 62, 65-67) and the MN (65). In addition, as one of the three deep cerebellar nuclei, the MN receives sensory information via vestibular nuclei (68) and projects to migraine-related brain regions including the thalamus (69). The MN can also be activated by noxious thermal stimuli (27). Moreover, injection of an excitatory amino acid into the MN decreased pain-related responses to visceral stimuli (70, 71). The same amino acid stimulation increased dorsal column nuclei activity in response to non-noxious somatic stimuli (72). These findings suggest that the MN, specifically CGRP receptors in the MN, may be associated with migraine pathophysiology. Thus, we hypothesized that CGRP injection into the MN might induce migraine-like behaviors in mice.

To address the role of cerebellar CGRP in migraine-like behaviors, we injected CGRP into the cerebellum centered on the MN and performed a battery of migraine-related behavioral tests. The results demonstrated that CGRP infusion into the MN induced light aversion in both sexes, while anxiety, tactile hypersensitivity, and squinting behaviors were predominately in female mice.

## 2 Materials and Methods

### 2.1 Animals

Wild-type C57BL/6J mice were obtained from Jackson Labs (Bar Harbor, ME and Sacramento, CA) at 8-12 weeks of age and were housed in groups of 2-5 per cage before surgery. A total of 55 C57BL/6J mice (28 females; 27 males) were used for this study. Female mice had an average starting body weight of 18-22 g and males were 20-25 g. Mice with cannulas were housed individually unless otherwise indicated to prevent mice from losing cannulas. All animals were housed on a 12-hour light cycle with access to water and food *ad libitum*. Animal procedures were approved by the Iowa City Veterans Administration and University of Iowa Animal Care and Use Committees and performed in accordance with the standards set by the National Institutes of Health.

### 2.2 Surgery

Cannulas were hand constructed from stainless steel hypodermic tubing (New England Small Tube Corporation; Supplementary Fig. 1). An 8-mm guide cannula was made from a 23-gauge needle (BD PrecisionGlide(tm)) with the ventral portion covered by a ∼7-mm, 19-gauge tubing (Supplementary Fig. 1A). The ∼7-mm, 19-gauge tubing is ∼2 mm higher than the 23-gauge needle to shield the junction between the guide’s top and the dummy or injection cannula after their insertion (Supplementary Fig. 1A). The dummy cannula, used to seal and keep the guide cannula free of clogs, was made by crimping a short segment of ∼5-mm, 23-gauge tubing to a ∼14 mm piece of 30-gauge tubing (Supplementary Fig. 1B). The bottom of the 30-gauge tubing was cut to ensure that the 30-gauge segment below the ∼5-mm, 23-gauge segment is 9 mm. The injection cannula was made by adhering a short segment of ∼5-mm, 23-gauge tubing ∼5 mm below the top of a ∼20-mm piece of 30-gauge tubing with adhesive (Pacer Technology) and dental cement (Stoelting) (Supplementary Fig. 1C). The bottom of the 30-gauge tubing was cut to ensure that the 30-gauge segment below the ∼5-mm, 23-gauge segment is 10 mm. In this manner, the dummy cannula extended 1 mm beyond the end of the guide cannula tip when it was inserted into the guide cannula, while the injection cannula protruded 2 mm from the base of the guide cannula (Supplementary Fig. 1D).

Stereotaxic implantation of a guide cannula into the MN of the right cerebellum was performed under isoflurane anesthesia (induction 5%, maintenance 1.5%–2%). The coordinates for the right MN are: anterior/posterior (AP), -6.5 mm posterior to bregma; medial/lateral (ML), -0.85 mm lateral to the midline; and dorsal/ventral (DV), -2.7 mm ventral to the pial surface according to the Allen Brain Reference Atlas. Guide cannulas were implanted 2 mm above the MN (AP: -6.5 mm; ML; -0.8 mm; DV: -0.7 mm). The implants were secured with bone anchor screws (Stoelting), adhesive, and dental cement. Dummy cannulas were inserted into guide cannulas when no injection was conducted. After surgery, mice were housed individually to reduce the loss of the guide or dummy cannulas. Mice were given ∼10 days to recover from the surgery before testing unless otherwise indicated.

### 2.3 Drug administration

Rat α-CGRP (Sigma-Aldrich) was diluted in 1X phosphate-buffered saline (PBS; HyClone^™^). Mice were given either rat α-CGRP (1 μg, 5 μg/μl) or 1X PBS (200 nl) as the vehicle through injection cannulas under anesthetized status (isoflurane: induction 5%, maintenance 1.5%–2%) unless otherwise indicated (details in Section 2.4.3 Von Frey test). Specifically, the dummy cannula was removed, and an injection cannula (2 mm extension from the base of the guide cannula) was inserted into the guide cannula. The injection cannula was connected to a 10 μl syringe (Hamilton) and an injection pump (Cole-Parmer Instrument Co.) via polyethylene tubing (BD Intramedic(tm), PE10). The injection rate was 100 nl/min for 2 min. After completing an infusion, the injection cannula was left in position for an additional 5-7 min before being withdrawn. Next, mice were returned to their home cages (individual housing) to recover for 60 min before testing unless otherwise indicated (details in Section 2.4.3 Von Frey test). The 60-min recovering period was chosen to minimize anesthesia effects (14, 15).

### 2.4 Behavioral tests

#### 2.4.1 Light/dark assay

The testing chamber was a transparent, seamless open field chamber divided into two zones of equal size by a black infrared-transparent dark insert (Med Associates). The mouse activity was collected with infrared beam tracking and Activity Monitor software (Med Associates), as previously described (14-16, 18, 73-76). Mice were tested without pre-exposure to the chamber using dim light (55 lux) after PBS or CGRP administration as described above. One hour post-injection, mice were placed in the light zone of the light/dark chamber and data were collected for 30 min and analyzed in sequential 5 min intervals.

Motility outcomes were collected during the light/dark assay, as described previously (14-16, 18, 73-76). Briefly, resting time was measured as the percentage of time animals did not break any new beams in each zone over the time spent in the same zone. Vertical beam breaks, an assessment of rearing behavior, was determined as the number of mice breaking the beam at 7.3-cm height in each zone, which was then normalized to the time spent in the same zone.

#### 2.4.2 Open field assay

This assay is to measure locomotion and anxiety-like behavior. The apparatus was the same as in the light/dark assay with the absence of the dark insert, as described previously (14, 18, 76). Mice were placed in the middle of the open field chamber with the light intensity at 55 lux one hour after PBS or CGRP infusion. The periphery was defined as 3.97 cm from the border with the reminding area of 19.05 × 19.05 cm as the center. The time in the center was calculated as the percentage of time spent in the center over the total time in the chamber.

#### 2.4.3 Von Frey test

The test is to evaluate the mechanical nociceptive threshold. For baseline experiments, mice were habituated to the room for one hour before acclimating to an acrylic chamber (10.80 × 6.99 × 14.61 cm in W x D x H) for one hour. The acrylic chamber was placed over a grid support (Bioseb, France). On the treatment day, investigators gently restrained the mouse and replaced the dummy cannula with an injection cannula. Then CGRP or PBS was infused via injection cannulas to the MN of the conscious and free-moving mice. Anesthesia (isoflurane) was not used since it induced a noticeable increase in the right hind paw withdrawal sensitivity in our pilot test. The reason is unclear, but one study reported that different doses of isoflurane had opposite effects on pain withdrawal sensitivity in response to thermal stimuli (77). Considering that the isoflurane effect might mask the CGRP effect, we decided to inject mice without anesthesia in the von Frey test. After injection, mice were allowed to rest in their home cages for 30 min and then placed in the acrylic chamber for another 30 min before applying von Frey filaments to their hind paws. Right and left hind paws were tested at the same time after treatment.

The investigator who applied filaments was blinded to the treatments and used the up-and-down method as previously described (78-80). Briefly, filaments were applied for 5 seconds to the skin of the mouse plantar surface, with D (0.07 g) as the starting filament. A withdrawal response was considered when mice withdrew, shook, or licked the tested hind paws. The withdrawal threshold at which 50% of mice withdrew their hind paws was determined based on an established equation (79, 80). However, the threshold data produced in this method are not continuous and cannot be analyzed using parametric statistics. Thus, in order to obtain normal distribution, the 50% thresholds (g) were transformed into log format for data analysis and figure plotting.

#### 2.4.4 Automated squint assay

This assay is to evaluate spontaneous pain by measuring the right-eye pixel areas recorded by a camera. Mice were acclimated to a customized collar restraint to reduce stress induced by restraint as well as struggle or head movement as described previously (81). C57BL/6J mice underwent acclimation for 20 min per session for three sessions. On the test day, after habituation to the room for one hour, the mouse was placed in the restraint, and squint was recorded for 5 min under room light as the baseline. Then CGRP or PBS was infused into the MN via an injection cannula. The mouse was returned to the home cage to rest for one hour, followed by another restraint and squint recording for 5 min under room light as the treatment recording. Pixel area measurement for the right eye palpebral fissure was derived every 0.1 seconds (10 frames per second) in the recordings using trained facial detection software (FaceX, LLC, Iowa City, IA) with the resulting values compiled with a custom MATLAB script. Individual frames containing a tracking error rate of >15% were excluded.

#### 2.4.5 Gait dynamic assay

Gait dynamics were measured using the DigiGait imaging system (Mouse Specifics Inc, Boston, MA, USA). The system consists of a transparent chamber (17.14 × 5.08 × 15.24 cm in W x D x H), a transparent plastic treadmill belt, an under-mounted digital camera, a light over the chamber for camera capturing videos (∼7200 lux), software to record videos (DigiGait Imager), and an image analysis software (DigiGait Analysis).

Mice were habituated to the room for one hour prior to any running. Mice first were placed in the chamber of the DigiGait apparatus for 1 min to allow them to explore the chamber. The belt was then turned on and mice were run at 16 cm/s, an optimal speed predetermined in C57BL/6J mice. Images of the paws were ventrally captured during the run. Each mouse ran until roughly 3-5 seconds of continuous gait was observed, a range sufficient to acquire adequate quantification of gait parameters. Mice underwent recordings before PBS or CGRP injection as the baseline. After injection, mice recovered in the home cages for one hour prior to another recording. A minimum of a one-hour interval was allotted between baseline and treatment trials to allow mice to recover from the previous running.

The mouse paw prints were analyzed by DigiGait Analysis to identify stride length and frequency. A complete stride was defined as the portion of foot strike to subsequent foot strike on the treadmill belt of the same foot.

### 2.5 Histology

After finishing all the behavioral tests, the injection sites were identified by the injection cannula tip, or by infusing Evans blue dye (200 nl, 1% dye, diluted in 1X PBS), or red retrograde beads (200 nl, Red Retrobeads^™^, LumaFluor, Inc.) to confirm targeting accuracy. Fluorescein-15-CGRP (1 μg; 200 nl mixed in 1X PBS) was injected into 4 mice to determine how far CGRP could spread from the MN. One hour post-injection, mice were deeply anesthetized with ketamine/xylazine (87.5 mg/kg/12.5 mg/kg, i.p.) and were perfused transcardially with 1X PBS and subsequently with 4% paraformaldehyde. Brains were removed and post-fixed in 4% paraformaldehyde at 4 °C overnight, followed by soaking in 10, 20, 30% sucrose per 24 hours in order. Brains were embedded in a tissue-freezing medium and stored at −80 °C until use. 100 μm coronal slices were collected from mouse brains injected with Evans blue. 40 μm coronal slices were collected from brains injected with red beads or fluorescein-15-CGRP. Slices from brains injected with fluorescein-15-CGRP were counterstained by incubation with TO-PRO-3 iodide. Slices were mounted onto Superfrost Plus slides (Fisher Scientific) using antifade mountant (VECTASHIELD). Images were captured using a scanning microscope (Olympus, VS120). Imaging of brains injected with Evans blue or red beads was performed using a light microscope (Olympus, CKX41) equipped with an Infinity 1 camera and processed using the INFINITY ANALYZE software (Lumenera Corporation).

### 2.6 Experimental design

To reduce the number of animals used in this study, the cannula system was used to allow the same mouse to undergo different assays. The first cohort was tested in the light/dark assay, open field assay, von Frey test, and automated squint assay. Because of the COVID-19 pandemic, the second cohort which had been exposed to the light/dark assay, open field assay and von Frey test were euthanized to minimize the burden in the animal facility. To repeat experiments in the automated squint assay, a third cohort was included. Unlike the previous two cohorts, the third cohort was first housed in groups after surgery. However, due to the high rate of dummy cannula loss in group housing conditions for about one week, mice were then housed individually instead, consistent with earlier cohorts. Von Frey test, gait dynamic and automated squint assays were performed in this cohort. Data from all cohorts were pooled for the final analysis.

The order of light/dark assay and open field assay was switched in the two cohorts to avoid an order bias. All the mice received the same treatment in the light/dark and open field assays to ensure the consistency. The same treatment in the light/dark assay was also given in the squint and gait dynamic assays. One cohort received crossover treatment in the automated squint assay. To ensure the withdrawal threshold in the von Frey test was comparable in control and experimental groups, mice were divided into two groups using a randomization protocol based on the baseline threshold. Mice were allowed to recover in their home cages for at least one day between each behavioral test. The light/dark or open field assays were performed first, followed by the von Frey test and the gait dynamic assay. The automated squint assay was performed last. All behavioral experiments were performed between 7:00 A.M. and 6:00 P.M, and mice were habituated to the room for one hour before experiments.

### 2.7 Statistical analysis

A power analysis was performed prior to experiments for sample size estimation based on previous studies from the lab and a post-hoc power analysis was performed to estimate the number of additional male mice needed to reach significance using ClinCalc.com. In the power analysis, an alpha of 0.05 and a power of 0.80 was used. The analysis determined that 10 mice in each group were needed. Data were analyzed using GraphPad Prism 9 and are reported in Supplementary Table 1.

Significance was set at P < 0.05. Error bars represent ± SEM. A two-way repeated measure ANOVA was performed when data were plotted as a function of time (factor: treatment and time) or for the scatter plot graphs of the von Frey and squint experiments (factor: treatment and condition). For all graphs, when the interaction or the condition was significant, Šídák’s multiple comparisons test was used as the post hoc analysis. For the data from the light/dark and open field assays, an unpaired t-test was performed for bar graphs with scatter points to compare the effect of each treatment.

A total of 3 mice died during the surgical procedure and one mouse lost the guide cannula before running any behavioral test. Two mice from the light/dark assay and one mouse from the open field assay were excluded due to chamber recording problems. In the von Frey test, 4 total mice were excluded: 3 mice due to the blockage of injection cannulas and one mouse due to the loss of the guide cannula. In the gait dynamic assay, 2 mice were excluded due to a video recording problem. In the automated squint assay, 2 mice were excluded due to the loss of the guide cannulas, 3 mice due to the blockage of injection cannulas, and 5 mice due to the poor habituation in the restraint or poor eye recognition by the software. Mouse numbers used for each experiment are reported in the figure legends.

## 3 Results

### 3.1 Injection of CGRP into the MN induced light-aversive behavior and reduced motility under dim light in both male and female mice

We injected CGRP into the MN of the right cerebellum via permanently placed cannulas and exposed the mice to the light/dark assay in dim light (55 lux) one hour post-injection. Light aversion was expressed as both a function of time over the 30-min testing period (Fig. 1A) and the average time in light for individual mice per 5 min interval (Fig. 1B). Regardless of sex, mice injected with CGRP spent less time in the light than those injected with PBS during the 30-min testing time (Fig. 1A, left). On average, the PBS-treated mice spent 141 seconds in the light per 5-min interval, and CGRP-treated mice spent 55 seconds (Fig.1B, left). When data were separated by sex, both male and female mice spent significantly less time in light after CGRP injection than those with PBS injection (Fig. 1A and B, middle and right). For all mice, confirmation of the targeting site was performed. The injection sites for mice in all behavioral tests are shown in Fig. 1C. Among 21 mice that experienced the light/dark assay, injection sites for 6 mice were not in or near the MN, primarily in the cerebellar cortex. However, these 6 off-target mice did not display significant differences in time in light from the on-target mice (data not shown). However, it should be noted that it is underpowered for off-target mice. Together, these data demonstrate that CGRP injection into the MN induces light-aversive behavior in both male and female mice.

**Fig. 1.**
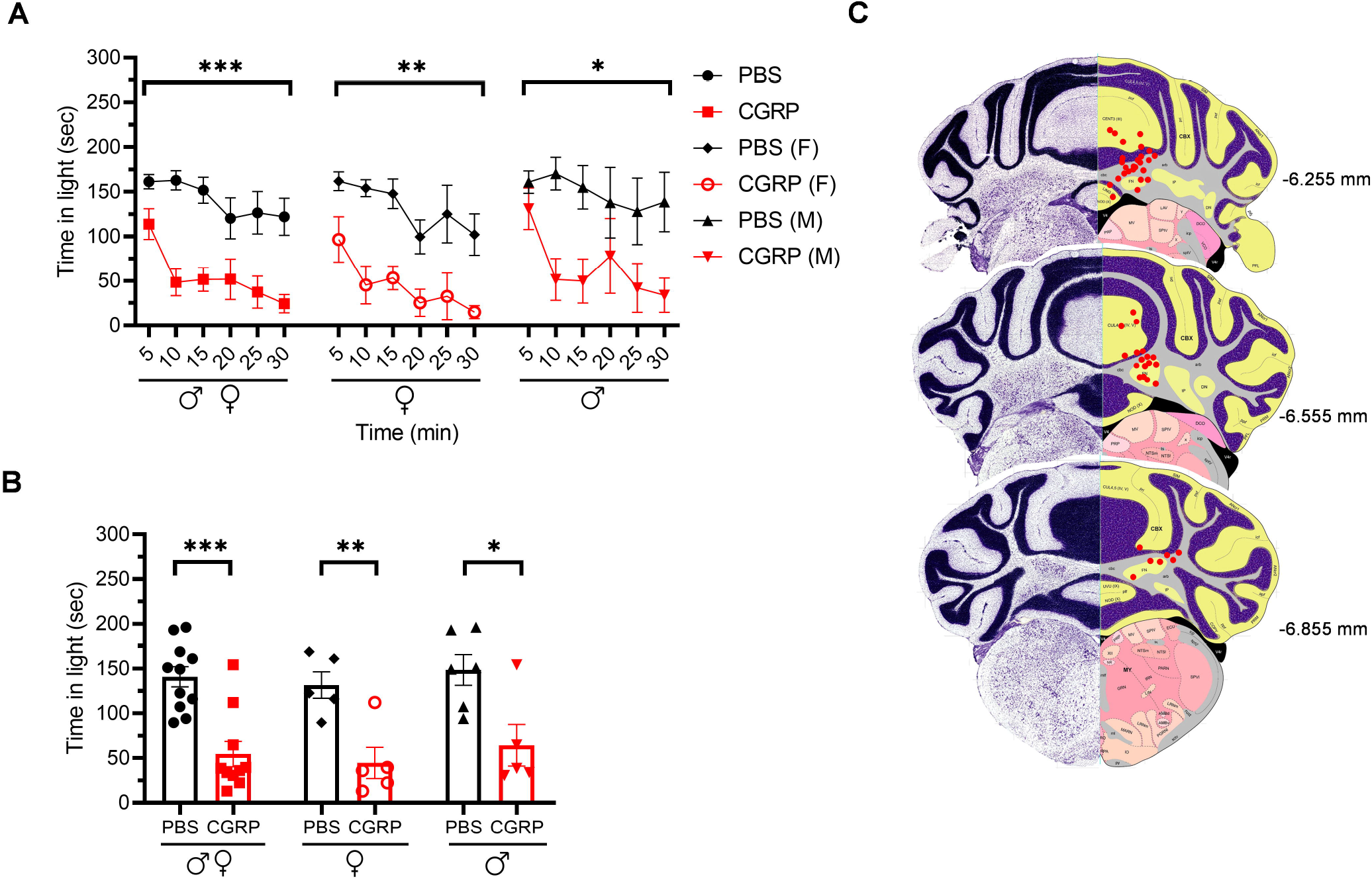
Injection of CGRP into the MN induced light-aversive behavior under dim light in both male and female mice. **(A)** Time in light every 5-min block during 30-min light/dark assay at 55 lux following injection of PBS (n=11; F: n=5; M; n=6) or CGRP (1 μg /200 nl; n=10; F: n=5; M: n=5) into the right MN of C57BL/6J mice via cannulas. Time in light for all mice (left), female mice (middle), and male mice (right). Data are from two independent experiments. All mice in A are further analyzed in B. **(B)** Mean time in light per 5-min block for individual mice. (**C**) Schematic of positions of injection cannula tips superimposed on Allen Mouse Brain Atlas coronal images. Numbers indicate the distance from bregma in the anteroposterior plane. Data are the mean ± SEM. Statistics are described in Supplementary Table 1.

Resting behavior was evaluated in the same light/dark assay. No difference was observed in the percent resting time in the light zone between CGRP- and PBS-injected mice (Fig. 2A and B, upper panel). In contrast, in the dark zone, CGRP-injected mice spent more time resting than PBS-injected mice across both sexes (Fig. 2A and B lower panel). In addition, CGRP-injected mice had significantly fewer rearings (vertical beams breaks) in both the light and dark zones across sexes (Fig. 2C and D). While there was a trend, the decreased rearing did not reach statistical significance in the male or female groups after CGRP injection, likely due to the variability and small sample size in each sex (Fig. 2C and D). Finally, transitions between dark and light zones were significantly decreased by CGRP for both sexes (Fig. 2E and F).

**Fig. 2.**
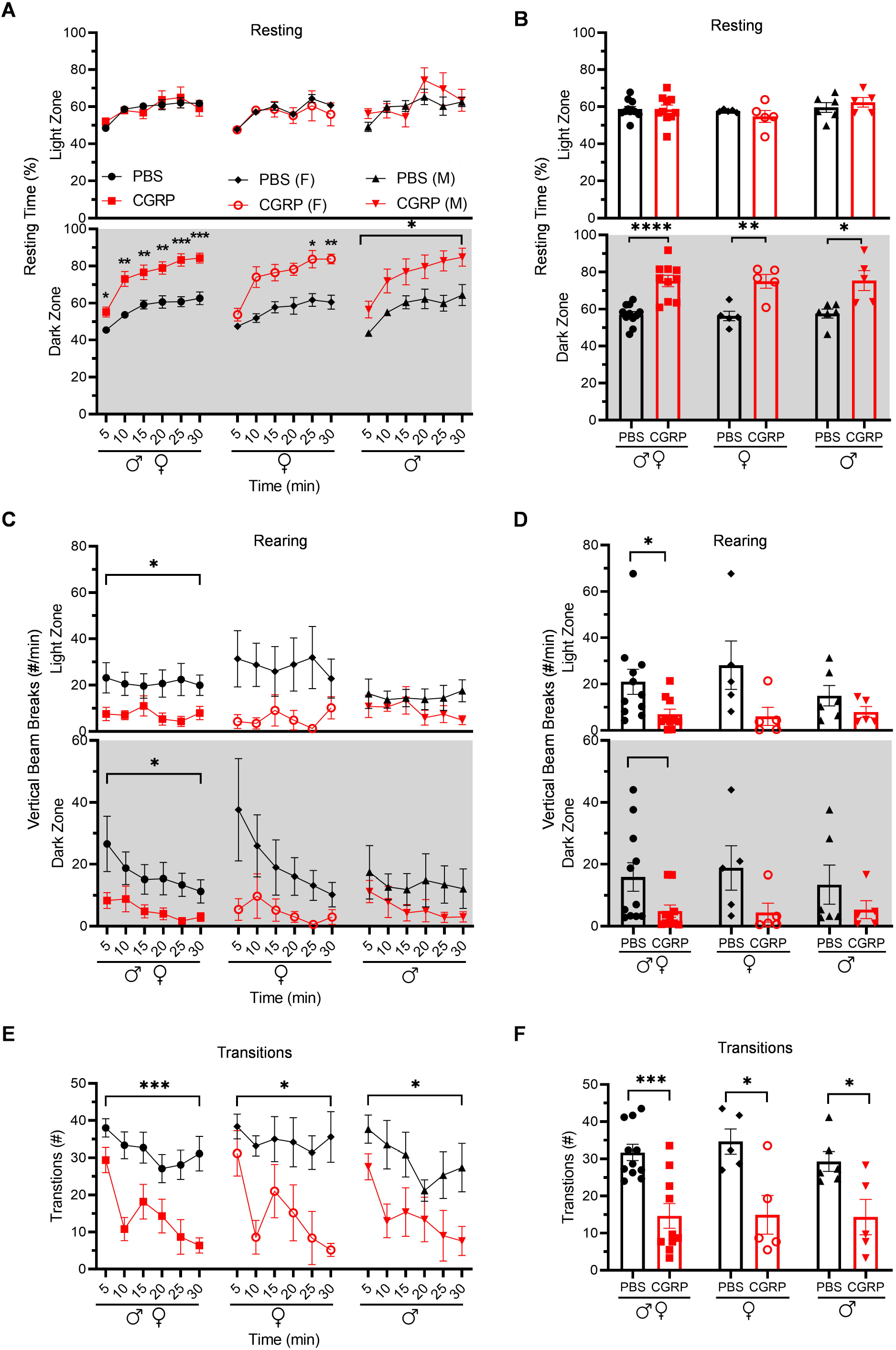
Injection of CGRP into the MN reduced motility in both males and females. Motility data were collected at the same time as light aversion data from the same mice shown in Fig. 1. Mice were given PBS (n=11; F: n=5; M: n=6) or CGRP (1 μg/200 nl; n=10; F: n=5; M: n=5) into the right MN of C57BL/6J mice via cannulas. Data are from two independent experiments. **(A)** Percentage of time spent resting in the light (upper panel) and dark (lower panel) zones every 5-min block during 30-min light/dark assay for all mice (left), female mice (middle), and male mice (right). All mice in A are further analyzed in B. **(B)** Mean percentage of time in light (upper panel) and dark (lower panel) zones per 5-min block for individual mice from A. **(C)** Number of vertical beam breaks per min in light (upper panel) and dark (lower panel) zones every 5-min block during 30-min light/dark assay for all mice (left), female mice (middle), and male mice (right). All mice in C are further analyzed in D. **(D)** Mean number of vertical beam breaks in light (upper panel) and dark (lower panel) zones per 5-min block for individual mice from C. **(E)** Number of transitions between light and dark zones every 5-min block during 30-min light/dark assay for all mice (left), female mice (middle), and male mice (right). All mice in E are further analyzed in F. **(F)** Mean number of transitions between light and dark zones per 5-min block for individual mice from E. Data are the mean ± SEM. Statistics are described in Supplementary Table 1.

Since the cerebellum is well-known for motor control and the MN controls axial and trunk muscles and maintains posture and balance (68), we tested the effect of CGRP delivery into the MN on gait. We conducted the gait dynamic assay using DigiGait system. Injection of PBS or CGRP into the MN did not change the stride length or frequency compared to their respective baselines across and within sexes (Supplementary Fig. 2). This indicates that CGRP administration into the MN decreases motility without gait alterations.

### 3.2 Injection of CGRP into the MN induced anxiety-like behavior primarily in female mice in the open field assay

To assess anxiety behavior independent of light, we used the open field assay. Inclusion of this assay was necessary because spending less time in the light in the light/dark assay can be an indicator of an increased anxiety state (76), and not necessarily a specific aversion to light. It is important to note though that the two are not mutually exclusive since light aversion may include increased anxiety.

The mice injected with CGRP spent less time in the center than those injected with PBS during the 30-min testing time (Fig. 3A and B, left). However, when data were analyzed by sex, females exhibited significantly less time in the center over the entire testing time (Fig. 3A and B, middle), while there was only a trend in males observed in the last 10 min (Fig. 3A and B, right). These data suggest that CGRP delivery into the MN elicited general anxiety-like behavior primarily in female mice, which may have contributed to their light-aversive behavior.

**Fig. 3.**
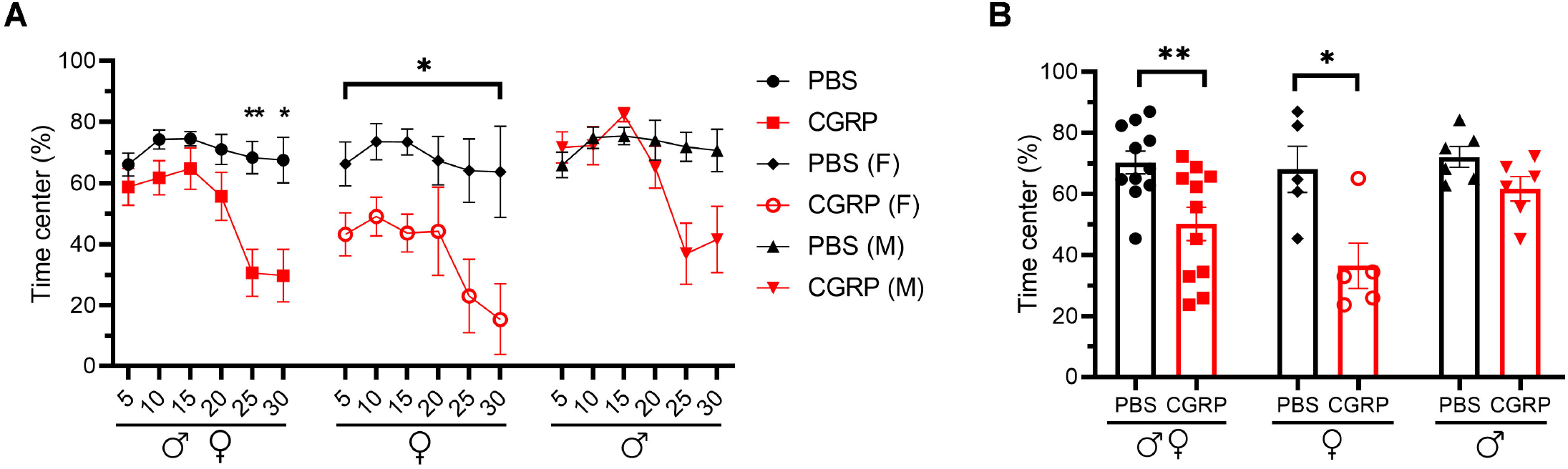
Injection of CGRP into the MN induced anxiety-like behavior primarily in females in the open field assay. **(A)** Percentage of time spent in the center of the open field every 5-min block during 30-min testing period following injection of PBS (n=11; F: n=5; M: n=6) or CGRP (1 μg/200 nl; n=11; F: n=5; M: n=6) into the right MN of C57BL/6J mice via cannulas. All mice (left) separated by sex (female: middle; male: right). Data are from two independent experiments. All mice in A are further analyzed in B. **(B)** Mean percentage of time in the center per 5-min block for individual mice. Data are the mean ± SEM. Statistics are described in Supplementary Table 1.

### 3.3 Injection of CGRP into the MN induced plantar tactile hypersensitivity in the contralateral hind paw primarily in female mice

Cutaneous allodynia is present in approximately 60% of migraine patients with a higher prevalence in women than men (82, 83). Thus, we investigated the effect of CGRP administration into the right MN on tactile hypersensitivity as a generally accepted indicator of allodynia by measuring the tactile sensitivity in the plantar area of the right and left hind paws.

In the contralateral left hind paw, there was a significant decrease in the withdrawal threshold observed for all the CGRP-treated mice (Fig. 4A, left). However, the difference is primarily driven by effects in the female mice, who showed a significant decrease in the withdrawal threshold after CGRP but not PBS injection compared to their respective baselines (Fig. 4A, middle). The CGRP-induced decrease in the threshold was only a trend in male mice (Fig. 4A, right).

**Fig. 4.**
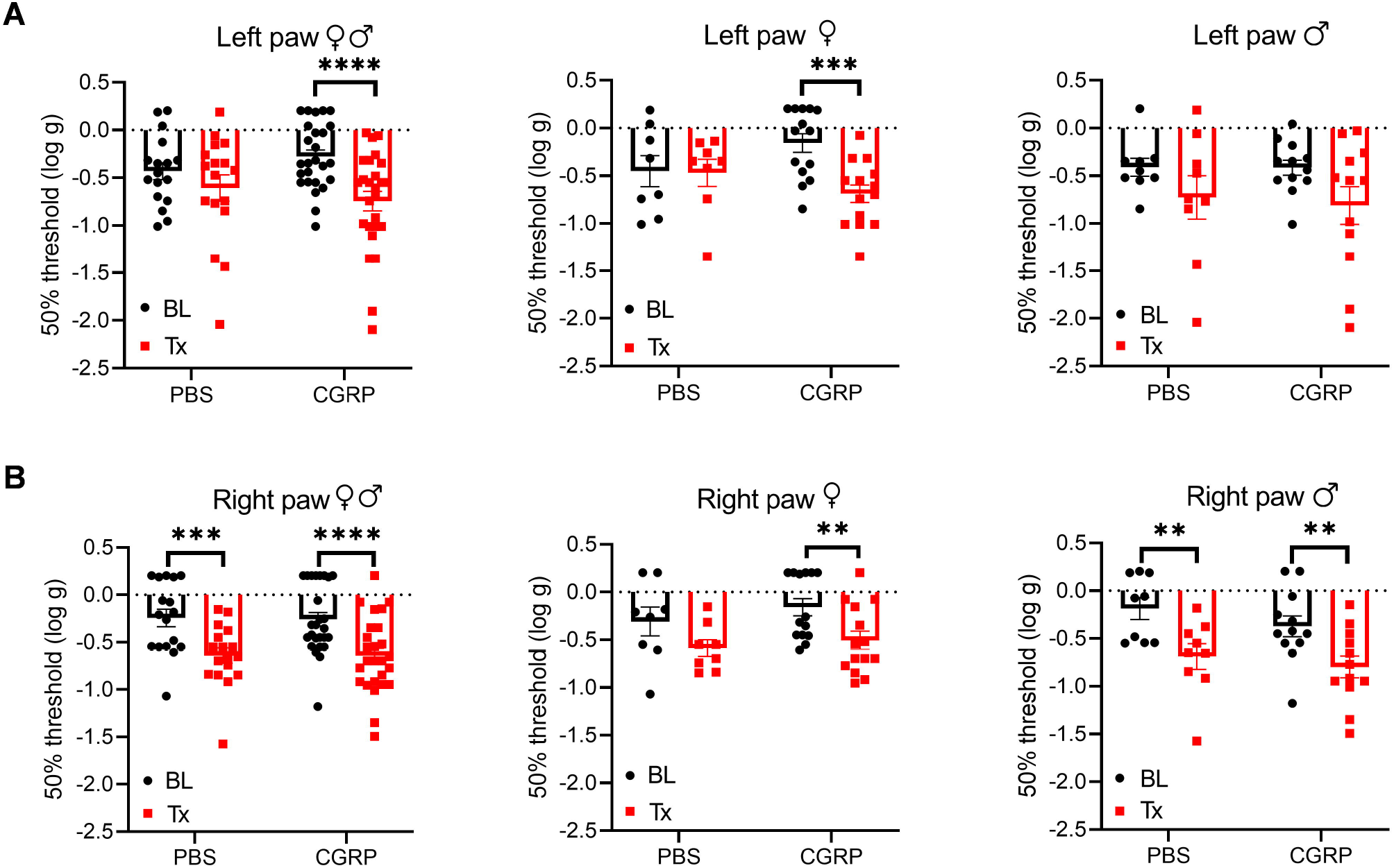
Injection of CGRP into the MN induced plantar tactile hypersensitivity in the contralateral hind paw primarily in female mice. Plantar tactile sensitivity was assessed with injection of PBS or CGRP (1 μg/200 nl) into the right MN of C57BL/6J mice via cannulas. Data are from three independent experiments. **(A)** The individual thresholds of left hind paws for all mice (left) (PBS: n=17; CGRP: n=26), female mice (middle) (PBS: n=8; CGRP: n=14), and male mice (right) (PBS: n=9; CGRP: n=12). **(B)** The individual thresholds of right hind paws for all mice (left) (PBS: n=17; CGRP: n=26), female mice (middle) (PBS: n=8; CGRP: n=14), and male mice (right) (PBS: n=9; CGRP: n=12). The mean ± SEM 50% thresholds are presented. Statistics are described in Supplementary Table 1.

In contrast, the ipsilateral right paw results were more complicated due to a significant decrease in withdrawal threshold compared to baselines in response to not only CGRP, but also PBS vehicle (Fig. 4B, left). When separated by sex, there was a trend for female mice after PBS treatment and a significant decrease after CGRP treatment compared to respective baselines (Fig. 4B, middle). A significant decrease was observed for male mice after either PBS or CGRP treatment (Fig. 4B, right) compared to baselines. The decrease in males after CGRP treatment is similar to the vehicle effect observed with PBS injection, suggesting that disturbance to the right MN is enough to increase ipsilateral hind paw sensitivity. For the von Frey test, 7 of the 43 mice had injection sites not in or near the MN. When comparing data between on-target mice and off-target mice, no difference was observed between these two groups, but it should be noted that the off-target mice were underpowered. Altogether, these data suggest that CGRP increases the contralateral left hind paw touch sensitivity predominantly in female mice, while injection of either PBS vehicle or CGRP increases sensitivity in the ipsilateral right hind paw.

### 3.4 Injection of CGRP into the MN induced squinting behavior primarily in female mice

The grimace scale was developed to evaluate spontaneous pain expression in mice (84). Our laboratory found that orbital tightening, or squint, is the principal component of mouse grimace score (85) and has developed an automated video-based squint assay to measure spontaneous pain (81). Taking advantage of this sensitive automated squint platform, we asked whether mice squint after CGRP injection in the MN.

For all mice, CGRP-treated mice showed a decrease in the mean pixel area over the 300-second testing period, while no change was observed in the PBS-treated mice compared to their respective baselines (Fig. 5A). When data were separated by sex, CGRP-treated females showed a significant decrease in the mean pixel area (Fig. 5B), while CGRP-treated males only showed a trend (Fig. 5C). No difference was observed in female or male PBS-treated groups (Fig. 5B and C, left and right).

**Fig. 5.**
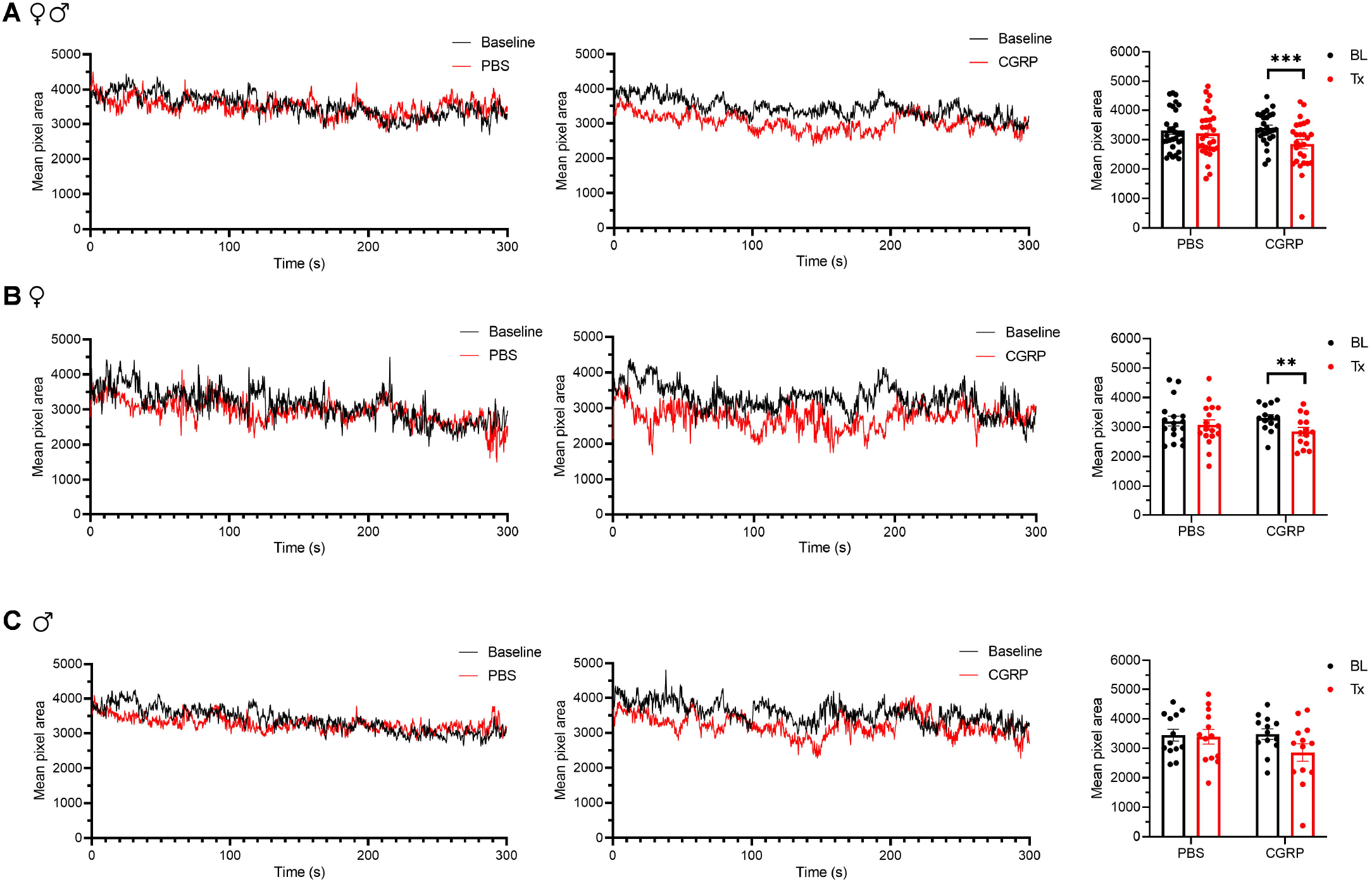
Injection of CGRP into the MN induced squinting behavior primarily in female mice. **(A)** Mean pixel area over 5-min testing period for all mice without treatment (as baseline), with injection of PBS (left) or CGRP (middle; 1 μg/200 nl) into the right MN of C57BL/6J mice via cannulas. Right panel is the mean pixel area over 5-min testing period for individual mice (PBS: n=30; CGRP: n= 27). Data from A separated as female (B) and male (C). Data are from two independent experiments and one crossover treatment experiment. **(B)** Mean pixel area over 5-min testing period for female mice (left and middle) and mean pixel area over 5-min testing period for individual female mice (right; PBS: n=17; CGRP: n= 14). **(C)** Mean pixel area over 5-min testing period for male mice (left and middle) and mean pixel area over 5-min testing period for individual male mice (right; PBS: n=13; CGRP: n=13). Data are the mean ± SEM. Statistics are described in Supplementary Table 1.

These data suggest that CGRP injection into the MN induces squint behavior predominantly in female mice.

### 3.5 The diffusion range of CGRP from the injection sites

To obtain an estimate of the likely diffusion of CGRP after injection into the MN, we used a fluorescent CGRP analog. Fluorescein-15-CGRP is a full CGRP receptor agonist but has less potency than CGRP as measured by cAMP production in HEK293T cells (86). Representative images of the rostral and caudal borders of fluorescein-15-CGRP diffusion from the MN are shown in Supplementary Fig. 3A upper and lower panels, respectively. There was considerable diffusion of fluorescein-15-CGRP from the injection site, with punctate signals found in cell bodies in the MN that may represent binding sites (Supplementary Fig. 3A, middle panel, box 1). In addition to the MN, fluorescein-15-CGRP was also observed in nearby regions, including the interposed and lateral cerebellar nuclei, granular, Purkinje cell, and molecular layers of vermal lobules I/III/IV/V (Supplementary Fig. 3A, middle panel, box 2). There was some variability in the spread of fluorescence among the mice injected with fluorescein-15-CGRP, with the smallest spread covering the MN and few nearby cells in the vermal lobules III/IV/V (Supplementary Fig. 3B, purple shading) and the largest spread covering the MN and cells beyond the MN including vermal lobules I/III/IV/V/X, the simple lobule and other cerebellar deep nuclei (Supplementary Fig. 3B, blue shading). The reason for variability in diffusion is not known but is apparently not due to injection site variability based on the injection sites shown by injection of red beads or Evans blue (Supplementary Fig. 3C and D).

## 4 Discussion

To our knowledge, this is the first preclinical cerebellar migraine study looking at behavioral outcomes. Indeed, there have been few animal studies looking at imaging and electrophysiological links between the cerebellum and migraine (87-92). Brain imaging studies have reported that the cerebellar sodium concentration and functional connectivity to the insula or anterior cingulate cortex were altered in animal migraine models induced by nitroglycerin or inflammatory soup (87-89). The firing rate of Purkinje cells in the rat paraflocculus was decreased in an animal model induced by trigeminal stimulation (90), and the organization of parallel fibers to Purkinje cell synapses was abnormal in familial hemiplegic migraine type 1 mouse models (91, 92). There are also preclinical behavioral studies investigating the role of the cerebellum in pain modulation (70-72, 93-99). Our finding that several migraine-like symptoms can be induced by CGRP actions in the cerebellum supports the hypothesis that the cerebellum contributes to migraine pathogenesis.

### 4.1 The MN and light aversion

Photophobia is a subjective experience in which normal light causes discomfort (100-102). In this study, we found that administration of CGRP into the MN induced light aversion in male and female mice. In addition, CGRP evoked anxiety-like behavior in females, but only a trend in males. Thus, we conclude that the light aversion induced in males is not solely driven by anxiety, while in females the light aversion may be influenced by an overall increased anxiety level. Anxiety symptoms have been reported to be positively correlated to light aversion in migraine patients with the possibility that anxiety contributes to light aversion (103). In male mice, there appears to be a biphasic response where an anxiety-like response may have occurred during the final 10 min of the assay. While not significant, it suggests that the light aversion detected in males may be partially driven by increased anxiety at later time points.

An unexpected finding was that both male and female mice were aversive to even dim light after CGRP injection, analogous to migraine patients who report light hypersensitivity in dim light that does not bother control subjects. We had previously reported that transgenic mice overexpressing the CGRP receptor in the nervous system were sensitive to dim light (∼55 lux) after i.c.v. CGRP injection (14, 16, 17), while light aversion induced in wild-type C57BL/6J mice required bright light (∼27,000 lux, similar to a sunny day) (15). Those data suggested that hypersensitivity to CGRP in the nervous system leads to light hypersensitivity. Interestingly, in contrast to i.c.v. injections, CGRP injected directly into the posterior thalamic nuclei (18) and cerebellar MN in this study, caused light aversion with dim light in C57BL/6J mice. These data indicate that like the posterior thalamus, the MN is sensitive to CGRP signaling without a need to increase receptor expression, perhaps due to increased local concentrations of CGRP relative to i.c.v. deliveries.

One model to explain the clinical manifestation of photophobia is convergence of signals from intrinsically photosensitive retinal ganglion cells onto posterior thalamic neurons that also receive nociceptive signals from the trigeminal nucleus (104). Light and nociceptive signals are then integrated and sent to the somatosensory and visual cortices (104). In support of this model, we have recently reported that injection of CGRP into the posterior thalamic region or optogenetic stimulation of that same region caused light aversion (18). How might the cerebellum fit into this model? One possibility may be via bilateral fibers from the principle sensory trigeminal nucleus and spinal trigeminal nucleus to the posterior vermis of the cerebellum (57), which projects to the MN (68). The MN is known to project to various thalamic nuclei including parafascicular, centrolateral, mediodorsal, ventrolateral, suprageniculate, and posterior nuclei (69). Thus, the MN may lie in a circuit from the trigeminal system to the thalamus. However, unlike the thalamus, there are no apparent direct retinocerebellar connections (105, 106). These data place the MN in a prime position to assist in sensory integration and play a modulatory role in the nociceptive- and light- integrating function of the thalamus.

### 4.2 The MN and anxiety

It is striking to observe the apparent sexually dimorphic anxiety-like behaviors predominantly in female mice after CGRP injection into the MN. These data are consistent with the higher prevalence of anxiety disorders in women than men (107). The MN sends direct projections to the limbic system including the amygdala (68), which is key to the anxiety circuitry (108), and projects to the periaqueductal gray (PAG) (69, 109), which is critical for aversive and anxiety-like responses (110). The observations of light-aversive behavior accompanied by increased anxiety levels in females are reminiscent of the behavior induced by optical stimulation of the dorsal PAG (18). This evidence might explain the anxiogenic effect of the MN. The possible mechanism for the sex difference in anxiety (or the other behaviors, including plantar tactile hypersensitivity and squinting discussed in Section 4.3) is not known. It is interesting to point out that sex differences appeared in human fMRI studies when migraine patients were exposed to a noxious stimulus (111). In that study, female migraine patients showed higher activation in the cerebellum and higher deactivation of cerebellar functional connectivity with insula than males in response to noxious heat. Finally, it is possible that there could be sexually dimorphic differences in the distribution pattern of CGRP receptor components in the MN or in downstream brain regions.

### 4.3 The MN and evoked and spontaneous pain

Allodynia is the perception of pain induced by non-noxious stimuli. Nearly 60% of individuals with migraine have cutaneous allodynia, specifically thermal and mechanical allodynia (82). Cutaneous allodynia is associated with migraine frequency, severity, and disability, and is more common in females (82, 83). Moreover, cutaneous allodynia is more frequent in chronic migraine than episodic migraine (112) and is believed to be a predictor of migraine chronification (113). Allodynia in migraine is found in cephalic and extracephalic regions, which could be explained by the sensitization of the second-order trigeminal and third-order thalamic neurons (114).

In this study, we found that CGRP injection into the MN increases sensitivity in response to mechanical stimuli in contralateral hind paws predominately in female mice, which is consistent with the clinical finding that cutaneous allodynia is higher in women than in men (82, 83) and a preclinical study where intraplantar CGRP at a low dose evoked hind paw allodynia only in female rats (115).

Given that we also observed anxiety behavior primarily in female mice, it is interesting that allodynia is associated with a higher risk for anxiety and a correlation exists between their severity in migraine patients (116). Anxiety was more prevalent in patients with migraine and probable migraine with cutaneous allodynia than those without cutaneous allodynia (117). In animal models, stress elicited higher pain sensitivity (118). These data suggest an association between anxiety and allodynia, so it is possible that anxiety induced by CGRP injection in female mice is linked to the tactile hypersensitivity indicative of allodynia.

How might the cerebellum increase paw sensitivity? There is evidence the cerebellum can affect the descending pain modulation pathway (93, 98, 99, 119), including via connections to the reticular formation (69, 93, 98). One study suggested that the MN could impact the dorsal column–medial lemniscus pathway directly or via the descending pain pathway (72). In addition, the MN projects to the thalamus bilaterally with contralateral preponderance (69), which might contribute to central sensitization and then lead to a pain hypersensitive state. However, the specific neuronal type in the MN that expresses CGRP receptors and specific regions that are modulated by CGRP in the MN are unknown.

An unexplained observation is that the ipsilateral hind paw showed a significant decrease in sensitivity after both PBS and CGRP injection into the MN. No change was observed after inserting the injection cannulas into the MN, without any injections, suggesting that the response was due to the solution. Because of the vehicle effect, a conclusion cannot be drawn from the ipsilateral paw data.

Our studies showed that CGRP injection into the MN induced squinting behavior predominately in females, suggesting CGRP in the MN plays a role in spontaneous pain. This is consistent with dural application of CGRP also causing grimace only in female mice (115). The magnitude of the squint response is relatively small in females (12.9%) compared to intraplantar injection of formalin (25.3%), and is closer to the response seen with wild-type female C57BL/6J mice receiving a small i.p. CGRP dose (0.01 mg/kg) (17.1%) (81).

### 4.4 The MN and motor function

We observed increased resting time in the dark while no change in the light in the light/dark assay across and within sex, corresponding to the preference of migraine patients to go to the dark and rest. Vertical beam breaks and transitions were decreased, suggesting exploratory behavior was decreased. The MN is responsible for controlling axial and trunk muscles, posture and balance (68). However, we did not observe gait difference before or after PBS or CGRP treatment using the DigiGait system. A recent study reported that an increase in light intensity could enhance postural sway in migraine patients compared to controls (54), so perhaps additional triggers may be needed to detect such effects in mouse models. Overall, these data suggest that CGRP injection into the MN does not induce gait changes.

### 4.5 Caveats

A caveat of this study is the broad diffusion area of CGRP. We used fluorescein-15-CGRP for diffusion estimation. Extensive spread was observed from the MN with the injection of fluorescein-15-CGRP, which was not completely unexpected. The diffusion was similar (approximately 800-4000 μm rostral to caudal) as when fluorescein-15-CGRP was injected into the posterior thalamic region and was estimated to spread at least 1400 um in some cases (18). This is consistent with the volume transmission reported for some peptides diffusing up to millimeters in the brain (120). The robust spread of CGRP explains why even the off-target injections had similar behaviors as the on-target injections. While the diffusion is extensive, it apparently did not reach the fourth ventricle, which is near the MN, since no fluorescein-15-CGRP was detected in the fourth ventricle.

Furthermore, the behavior is not likely due to CGRP diffusing into the cerebrospinal fluid since, as mentioned earlier, i.c.v. CGRP did not induce light aversion in wild-type C57BL/6J mice under dim light (15), while injection into the MN did. Nonetheless, the broad diffusion of fluorescein-15-CGRP beyond the MN decreases the targeting specificity, which makes it difficult to pinpoint which regions, in the MN or near the MN, are important for the responses induced by CGRP. Future studies will need to focus on limiting the spread of CGRP beyond the MN.

A related caveat is that location of CGRP receptor subunits RAMP1 and CLR in the mouse cerebellum has not been studied. Importantly, clusters of fluorescein-15-CGRP observed within the MN are consistent with a prior report of MN CGRP receptors in the rat (65). Such clusters were also found in the molecular, Purkinje cell and granular layers in the vermal lobules and simple lobule in the hemisphere regions. Previous studies have reported RAMP1 and CLR co-expression in Purkinje cells in the rat, human and rhesus cerebellum (61, 66). However, consistent data is lacking for RAMP1 or/and CLR expression in the molecular layer or granular layer in the rat cerebellum (61, 65) and no reports for mice, to our knowledge. In addition, the possible expression of the second CGRP receptor (AMY1) (121) in the mouse cerebellum has not been explored.

We observed several responses to cerebellar CGRP that were only statistically significant in female mice. However, we want to couch that observation with the prediction that if more mice were analyzed, then the trends seen with male mice could also reach significance for the open field, von Frey, and grimace assays. Because the study was designed to be sufficiently powered (see section 2.7), we did not try to further increase the number of male mice. However, to estimate how many more mice might be required to reach significance, we did a post-hoc power analysis for each of the assays. About twice the number of male mice is predicted to be needed to reach statistical significance comparable to the females. Hence, as seen with human migraine populations, we are seeing a quantitative bias for female responses and not an absolute female-only mechanism.

### 4.6 Conclusions

In conclusion, this study reveals that CGRP injection into the cerebellum is sufficient to induce migraine-like behaviors primarily in female mice. This discovery provides a new perspective on the increasingly complex neural circuitry of migraine pathophysiology and suggests a role for central CGRP signaling in the sexual dimorphic nature of migraine.

## 6 Author Contributions

Author contributions: M.W., L.P.S., and A.F.R. designed research; M.W., T.L.D., B.J.R., J.S.W., M.W.H., H.C.F. performed research; M.W., L.P.S., T.L.D. analyzed data; M.W., L.P.S., T.L.D., A.F.R. interpreted data; M.W., L.P.S., A.F.R. wrote the paper.

## 8 Acknowledgments

We thank Krystal Parker, Jonah Heskje and Hunter Halverson for the help on the cannula system, Debbie Hay and Christopher Walker for providing fluorescein-15-CGRP, and the VA Center for the Prevention and Treatment of Visual Loss for use of facilities.

## References

1. Collaborators GBDH. Global, regional, and national burden of migraine and tension-type headache, 1990-2016: a systematic analysis for the Global Burden of Disease Study 2016. Lancet Neurol (2018) 17(11):954–76. Epub 2018/10/26. doi: 10.1016/S1474-4422(18)30322-3. PubMed PMID: 30353868; PubMed Central PMCID: PMCPMC6191530.

2. Disease GBD, Injury I, Prevalence C. Global, regional, and national incidence, prevalence, and years lived with disability for 354 diseases and injuries for 195 countries and territories, 1990-2017: a systematic analysis for the Global Burden of Disease Study 2017. Lancet (2018) 392(10159):1789–858. Epub 2018/11/30. doi: 10.1016/S0140-6736(18)32279-7. PubMed PMID: 30496104; PubMed Central PMCID: PMCPMC6227754.

3. Headache Classification Committee of the International Headache Society (IHS) The International Classification of Headache Disorders, 3rd edition. Cephalalgia (2018) 38(1):1–211. Epub 2018/01/26. doi: 10.1177/0333102417738202. PubMed PMID: 29368949.

4. Goadsby PJ, Edvinsson L, Ekman R. Vasoactive peptide release in the extracerebral circulation of humans during migraine headache. Ann Neurol (1990) 28(2):183–7. Epub 1990/08/01. doi: 10.1002/ana.410280213. PubMed PMID: 1699472.

5. Ashina M, Bendtsen L, Jensen R, Schifter S, Olesen J. Evidence for increased plasma levels of calcitonin gene-related peptide in migraine outside of attacks. Pain (2000) 86(1-2):133–8. Epub 2000/04/26. doi: 10.1016/s0304-3959(00)00232-3. PubMed PMID: 10779670.

6. Cernuda-Morollon E, Larrosa D, Ramon C, Vega J, Martinez-Camblor P, Pascual J. Interictal increase of CGRP levels in peripheral blood as a biomarker for chronic migraine. Neurology (2013) 81(14):1191–6. doi: DOI 10.1212/WNL.0b013e3182a6cb72. PubMed PMID: WOS:000330768200007.

7. Asghar MS, Hansen AE, Amin FM, van der Geest RJ, Koning P, Larsson HB, et al. Evidence for a vascular factor in migraine. Ann Neurol (2011) 69(4):635–45. Epub 2011/03/19. doi: 10.1002/ana.22292. PubMed PMID: 21416486.

8. Guo S, Vollesen ALH, Olesen J, Ashina M. Premonitory and nonheadache symptoms induced by CGRP and PACAP38 in patients with migraine. PAIN (2016) 157(12):2773–81. doi: 10.1097/j.pain.0000000000000702. PubMed PMID: 00006396-201612000-00018.

9. Hansen JM, Hauge AW, Olesen J, Ashina M. Calcitonin gene-related peptide triggers migraine-like attacks in patients with migraine with aura. Cephalalgia (2010) 30(10):1179–86. doi: 10.1177/0333102410368444. PubMed PMID: WOS:000285052900005.

10. Lassen LH, Haderslev P, Jacobsen VB, Iversen HK, Sperling B, Olesen J. CGRP may play a causative role in migraine. Cephalalgia (2002) 22(1):54–61. doi: DOI 10.1046/j.1468-2982.2002.00310.x. PubMed PMID: WOS:000174216500009.

11. Edvinsson L, Haanes KA, Warfvinge K, Krause DN. CGRP as the target of new migraine therapies - successful translation from bench to clinic. Nat Rev Neurol (2018) 14(6):338–50. doi: 10.1038/s41582-018-0003-1. PubMed PMID: WOS:000433426600010.

12. Rapoport AM, McAllister P. The Headache Pipeline: Excitement and Uncertainty. Headache (2020) 60(1):190–9. doi: 10.1111/head.13728. PubMed PMID: WOS:000506067400020.

13. Johnson KW, Morin SM, Wroblewski VJ, Johnson MP. Peripheral and central nervous system distribution of the CGRP neutralizing antibody [I-125] galcanezumab in male rats. Cephalalgia (2019) 39(10):1241–8. doi: 10.1177/0333102419844711. PubMed PMID: WOS:000482243200005.

14. Mason BN, Kaiser EA, Kuburas A, Loomis MM, Latham JA, Garcia-Martinez LF, et al. Induction of Migraine-Like Photophobic Behavior in Mice by Both Peripheral and Central CGRP Mechanisms. J Neurosci (2017) 37(1):204–16. Epub 2017/01/06. doi: 10.1523/JNEUROSCI.2967-16.2016. PubMed PMID: 28053042; PubMed Central PMCID: PMCPMC5214631.

15. Kaiser EA, Kuburas A, Recober A, Russo AF. Modulation of CGRP-induced light aversion in wild-type mice by a 5-HT(1B/D) agonist. J Neurosci (2012) 32(44):15439–49. Epub 2012/11/02. doi: 10.1523/JNEUROSCI.3265-12.2012. PubMed PMID: 23115181; PubMed Central PMCID: PMCPMC3498941.

16. Recober A, Kaiser EA, Kuburas A, Russo AF. Induction of multiple photophobic behaviors in a transgenic mouse sensitized to CGRP. Neuropharmacology (2010) 58(1):156–65. Epub 2009/07/18. doi: 10.1016/j.neuropharm.2009.07.009. PubMed PMID: 19607849; PubMed Central PMCID: PMCPMC2784010.

17. Recober A, Kuburas A, Zhang Z, Wemmie JA, Anderson MG, Russo AF. Role of calcitonin gene-related peptide in light-aversive behavior: implications for migraine. J Neurosci (2009) 29(27):8798–804. Epub 2009/07/10. doi: 10.1523/JNEUROSCI.1727-09.2009. PubMed PMID: 19587287; PubMed Central PMCID: PMCPMC2944225.

18. Sowers LP, Wang M, Rea BJ, Taugher RJ, Kuburas A, Kim Y, et al. Stimulation of Posterior Thalamic Nuclei Induces Photophobic Behavior in Mice. Headache (2020) 60(9):1961–81. Epub 2020/08/05. doi: 10.1111/head.13917. PubMed PMID: 32750230; PubMed Central PMCID: PMCPMC7604789.

19. Rondi-Reig L, Paradis A-L, Lefort JM, Babayan BM, Tobin C. How the cerebellum may monitor sensory information for spatial representation. Front Syst Neurosci (2014) 8:205–. doi: 10.3389/fnsys.2014.00205. PubMed PMID: 25408638.

20. Wiestler T, McGonigle DJ, Diedrichsen J. Integration of sensory and motor representations of single fingers in the human cerebellum. J Neurophysiol (2011) 105(6):3042–53. Epub 2011/04/08. doi: 10.1152/jn.00106.2011. PubMed PMID: 21471398.

21. Manto M, Bower JM, Conforto AB, Delgado-Garcia JM, da Guarda SNF, Gerwig M, et al. Consensus Paper: Roles of the Cerebellum in Motor Control-The Diversity of Ideas on Cerebellar Involvement in Movement. Cerebellum (2012) 11(2):457–87. doi: 10.1007/s12311-011-0331-9. PubMed PMID: WOS:000304466100026.

22. Baumann O, Borra RJ, Bower JM, Cullen KE, Habas C, Ivry RB, et al. Consensus Paper: The Role of the Cerebellum in Perceptual Processes. Cerebellum (2015) 14(2):197–220. doi: 10.1007/s12311-014-0627-7. PubMed PMID: WOS:000350897800014.

23. Adamaszek M, D’Agata F, Ferrucci R, Habas C, Keulen S, Kirkby KC, et al. Consensus Paper: Cerebellum and Emotion. Cerebellum (2017) 16(2):552–76. doi: 10.1007/s12311-016-0815-8. PubMed PMID: WOS:000396044200024.

24. Van Overwalle F, Manto M, Cattaneo Z, Clausi S, Ferrari C, Gabrieli JDE, et al. Consensus Paper: Cerebellum and Social Cognition. Cerebellum (2020) 19(6):833–68. doi: 10.1007/s12311-020-01155-1. PubMed PMID: WOS:000545937300001.

25. Koziol LF, Budding D, Andreasen N, D’Arrigo S, Bulgheroni S, Imamizu H, et al. Consensus Paper: The Cerebellum’s Role in Movement and Cognition. Cerebellum (2014) 13(1):151–77. doi: 10.1007/s12311-013-0511-x. PubMed PMID: WOS:000332156100016.

26. Parker KL, Narayanan NS, Andreasen NC. The therapeutic potential of the cerebellum in schizophrenia. Front Syst Neurosci (2014) 8:163. Epub 2014/10/14. doi: 10.3389/fnsys.2014.00163. PubMed PMID: 25309350; PubMed Central PMCID: PMCPMC4163988.

27. Moulton EA, Schmahmann JD, Becerra L, Borsook D. The cerebellum and pain: passive integrator or active participator? Brain Res Rev (2010) 65(1):14–27. Epub 2010/06/18. doi: 10.1016/j.brainresrev.2010.05.005. PubMed PMID: 20553761; PubMed Central PMCID: PMCPMC2943015.

28. Farago P, Tuka B, Toth E, Szabo N, Kiraly A, Csete G, et al. Interictal brain activity differs in migraine with and without aura: resting state fMRI study. J Headache Pain (2017) 18. doi: ARTN 8 10.1186/s10194-016-0716-8. PubMed PMID: WOS:000397741200001.

29. Wang JJ, Chen X, Sah SK, Zeng C, Li YM, Li N, et al. Amplitude of low-frequency fluctuation (ALFF) and fractional ALFF in migraine patients: a resting-state functional MRI study. Clin Radiol (2016) 71(6):558–64. doi: 10.1016/j.crad.2016.03.004. PubMed PMID: WOS:000376697000010.

30. Afridi SK, Giffin NJ, Kaube H, Friston KJ, Ward NS, Frackowiak RSJ, et al. A positron emission tomographic study in spontaneous migraine. Arch Neurol-Chicago (2005) 62(8):1270–5. doi: DOI 10.1001/archneur.62.8.1270. PubMed PMID: WOS:000231034300015.

31. Bonanno L, Lo Buono V, De Salvo S, Ruvolo C, Torre V, Bramanti P, et al. Brain morphologic abnormalities in migraine patients: an observational study. J Headache Pain (2020) 21(1). doi: 10.1186/s10194-020-01109-2. PubMed PMID: WOS:000530354300003.

32. Demir BT, Bayram NA, Ayturk Z, Erdamar H, Seven P, Calp A, et al. Structural Changes in the Cerebrum, Cerebellum and Corpus Callosum in Migraine Patients. Clin Invest Med (2016) 39(6):S21–S6. PubMed PMID: WOS:000389725000005.

33. Jin CW, Yuan K, Zhao LM, Zhao L, Yu DH, von Deneen KM, et al. Structural and functional abnormalities in migraine patients without aura. Nmr Biomed (2013) 26(1):58–64. doi: 10.1002/nbm.2819. PubMed PMID: WOS:000313886200007.

34. Mehnert J, May A. Functional and structural alterations in the migraine cerebellum. J Cerebr Blood F Met (2019) 39(4):730–9. doi: 10.1177/0271678x17722109. PubMed PMID: WOS:000463032400012.

35. Yang FC, Chou KH, Lee PL, Yin JH, Chen SY, Kao HW, et al. Patterns of gray matter alterations in migraine and restless legs syndrome. Ann Clin Transl Neur (2019) 6(1):57–67. doi: 10.1002/acn3.680. PubMed PMID: WOS:000455915000006.

36. Bilgic B, Kocaman G, Arslan AB, Noyan H, Sherifov R, Alkan A, et al. Volumetric differences suggest involvement of cerebellum and brainstem in chronic migraine. Cephalalgia (2016) 36(4):301–8. doi: 10.1177/0333102415588328. PubMed PMID: WOS:000374014500001.

37. Coppola G, Petolicchio B, Di Renzo A, Tinelli E, Di Lorenzo C, Parisi V, et al. Cerebral gray matter volume in patients with chronic migraine: correlations with clinical features. J Headache Pain (2017) 18. doi: ARTN 115 10.1186/s10194-017-0825-z. PubMed PMID: WOS:000419913100001.

38. Lai TH, Chou KH, Fuh JL, Lee PL, Kung YC, Lin CP, et al. Gray matter changes related to medication overuse in patients with chronic migraine. Cephalalgia (2016) 36(14):1324–33. doi: 10.1177/0333102416630593. PubMed PMID: WOS:000390004500003.

39. Elliott MA, Peroutka SJ, Welch S, May EF. Familial hemiplegic migraine, nystagmus, and cerebellar atrophy. Ann Neurol (1996) 39(1):100–6. Epub 1996/01/01. doi: 10.1002/ana.410390115. PubMed PMID: 8572654.

40. Haan J, Terwindt GM, Bos PL, Ophoff RA, Frants RR, Ferrari MD. Familial hemiplegic migraine in The Netherlands. Dutch Migraine Genetics Research Group. Clin Neurol Neurosurg (1994) 96(3):244–9. Epub 1994/08/01. doi: 10.1016/0303-8467(94)90076-0. PubMed PMID: 7988094.

41. Joutel A, Bousser MG, Biousse V, Labauge P, Chabriat H, Nibbio A, et al. A gene for familial hemiplegic migraine maps to chromosome 19. Nat Genet (1993) 5(1):40–5. Epub 1993/09/01. doi: 10.1038/ng0993-40. PubMed PMID: 8220421.

42. Kono S, Terada T, Ouchi Y, Miyajima H. An altered GABA-A receptor function in spinocerebellar ataxia type 6 and familial hemiplegic migraine type 1 associated with the CACNA1A gene mutation. BBA Clin (2014) 2:56–61. Epub 2015/12/18. doi: 10.1016/j.bbacli.2014.09.005. PubMed PMID: 26675662; PubMed Central PMCID: PMCPMC4633947.

43. Vighetto A, Froment JC, Trillet M, Aimard G. Magnetic resonance imaging in familial paroxysmal ataxia. Arch Neurol (1988) 45(5):547–9. Epub 1988/05/01. doi: 10.1001/archneur.1988.00520290083018. PubMed PMID: 3358708.

44. Huang XB, Zhang D, Chen YC, Wang P, Mao CN, Miao ZF, et al. Altered functional connectivity of the red nucleus and substantia nigra in migraine without aura. J Headache Pain (2019) 20(1). doi: ARTN 104 10.1186/s10194-019-1058-0. PubMed PMID: WOS:000496421100002.

45. Ke J, Yu Y, Zhang XD, Su YY, Wang XM, Hu S, et al. Functional Alterations in the Posterior Insula and Cerebellum in Migraine Without Aura: A Resting-State MRI Study. Front Behav Neurosci (2020) 14. doi: ARTN 567588 10.3389/fnbeh.2020.567588. PubMed PMID: WOS:000579489100001.

46. Moulton EA, Becerra L, Johnson A, Burstein R, Borsook D. Altered Hypothalamic Functional Connectivity with Autonomic Circuits and the Locus Coeruleus in Migraine. Plos One (2014) 9(4). doi: ARTN e95508 10.1371/journal.pone.0095508. PubMed PMID: WOS:000335309100131.

47. Qin ZX, Su JJ, He XW, Ban SY, Zhu Q, Cui YY, et al. Disrupted functional connectivity between sub-regions in the sensorimotor areas and cortex in migraine without aura. J Headache Pain (2020) 21(1). doi: ARTN 47 10.1186/s10194-020-01118-1. PubMed PMID: WOS:000533907700003.

48. Wei HL, Chen JA, Chen YC, Yu YS, Zhou GP, Qu LJ, et al. Impaired functional connectivity of limbic system in migraine without aura. Brain Imaging Behav (2020) 14(5):1805–14. doi: 10.1007/s11682-019-00116-5. PubMed PMID: WOS:000579512100044.

49. Zhang D, Huang XB, Su W, Chen YC, Wang P, Mao CN, et al. Altered lateral geniculate nucleus functional connectivity in migraine without aura: a resting-state functional MRI study. J Headache Pain (2020) 21(1). doi: ARTN 17 10.1186/s10194-020-01086-6. PubMed PMID: WOS:000514509200001.

50. Ziegeler C, Mehnert J, Asmussen K, May A. Central effects of erenumab in migraine patients: An event-related functional imaging study. Neurology (2020) 95(20):e2794–e802. Epub 2020/09/13. doi: 10.1212/WNL.0000000000010740. PubMed PMID: 32917805.

51. Karatas M. Migraine and Vertigo. Headache Research and Treatment (2011) 2011:793672. doi: 10.1155/2011/793672.

52. Anagnostou E, Gerakoulis S, Voskou P, Kararizou E. Postural instability during attacks of migraine without aura. Eur J Neurol (2019) 26(2):319–+. doi: 10.1111/ene.13815. PubMed PMID: WOS:000455803800019.

53. Ishizaki K, Mori N, Takeshima T, Fukuhara Y, Ijiri T, Kusumi M, et al. Static stabilometry in patients with migraine and tension-type headache during a headache-free period. Psychiatry Clin Neurosci (2002) 56(1):85–90. Epub 2002/04/04. doi: 10.1046/j.1440-1819.2002.00933.x. PubMed PMID: 11929575.

54. Pinheiro CF, Moraes R, Carvalho GF, Sestari L, Will-Lemos T, Bigal ME, et al. The Influence of Photophobia on Postural Control in Patients With Migraine. Headache (2020) 60(8):1644–52. doi: 10.1111/head.13908. PubMed PMID: WOS:000558689000001.

55. Hayashi H, Sumino R, Sessle BJ. Functional organization of trigeminal subnucleus interpolaris: nociceptive and innocuous afferent inputs, projections to thalamus, cerebellum, and spinal cord, and descending modulation from periaqueductal gray. J Neurophysiol (1984) 51(5):890–905. Epub 1984/05/01. doi: 10.1152/jn.1984.51.5.890. PubMed PMID: 6726316.

56. Ohya A, Tsuruoka M, Imai E, Fukunaga H, Shinya A, Furuya R, et al. Thalamic-Projecting and Cerebellar-Projecting Interpolaris Neuron Responses to Afferent Inputs. Brain Res Bull (1993) 32(6):615–21. doi: DOI 10.1016/0361-9230(93)90163-6. PubMed PMID: WOS:A1993LX83100009.

57. Ge SN, Li ZH, Tang J, Ma Y, Hioki H, Zhang T, et al. Differential expression of VGLUT1 or VGLUT2 in the trigeminothalamic or trigeminocerebellar projection neurons in the rat. Brain Struct Funct (2014) 219(1):211–29. Epub 2013/02/06. doi: 10.1007/s00429-012-0495-1. PubMed PMID: 23380804.

58. Baldacara L, Borgio JG, Lacerda AL, Jackowski AP. Cerebellum and psychiatric disorders. Braz J Psychiatry (2008) 30(3):281–9. Epub 2008/10/04. doi: 10.1590/s1516-44462008000300016. PubMed PMID: 18833430.

59. Hostetler ED, Joshi AD, Sanabria-Bohorquez S, Fan H, Zeng Z, Purcell M, et al. In vivo quantification of calcitonin gene-related peptide receptor occupancy by telcagepant in rhesus monkey and human brain using the positron emission tomography tracer [11C]MK-4232. J Pharmacol Exp Ther (2013) 347(2):478–86. Epub 2013/08/27. doi: 10.1124/jpet.113.206458. PubMed PMID: 23975906.

60. Salvatore CA, Moore EL, Calamari A, Cook JJ, Michener MS, O’Malley S, et al. Pharmacological properties of MK-3207, a potent and orally active calcitonin gene-related peptide receptor antagonist. J Pharmacol Exp Ther (2010) 333(1):152–60. Epub 2010/01/13. doi: 10.1124/jpet.109.163816. PubMed PMID: 20065019.

61. Edvinsson L, Eftekhari S, Salvatore CA, Warfvinge K. Cerebellar distribution of calcitonin gene-related peptide (CGRP) and its receptor components calcitonin receptor-like receptor (CLR) and receptor activity modifying protein 1 (RAMP1) in rat. Mol Cell Neurosci (2011) 46(1):333–9. Epub 2010/11/03. doi: 10.1016/j.mcn.2010.10.005. PubMed PMID: 21040789.

62. Eftekhari S, Salvatore CA, Connolly BM, O’Malley S, Miller PJ, Zeng Z, et al. CGRP and CGRP receptors in human and rhesus monkey cerebellum. J Headache Pain (2013) 14. doi: Artn P91 10.1186/1129-2377-14-S1-P91. PubMed PMID: WOS:000325525600109.

63. Morara S, Rosina A, Provini L, Forloni G, Caretti A, Wimalawansa SJ. Calcitonin gene-related peptide receptor expression in the neurons and glia of developing rat cerebellum: an autoradiographic and immunohistochemical analysis. Neuroscience (2000) 100(2):381–91. Epub 2000/09/29. doi: 10.1016/s0306-4522(00)00276-1. PubMed PMID: 11008176.

64. Morara S, Wimalawansa SJ, Rosina A. Monoclonal antibodies reveal expression of the CGRP receptor in Purkinje cells, interneurons and astrocytes of rat cerebellar cortex. Neuroreport (1998) 9(16):3755–9. Epub 1998/12/19. doi: 10.1097/00001756-199811160-00034. PubMed PMID: 9858392.

65. Warfvinge K, Edvinsson L. Distribution of CGRP and CGRP receptor components in the rat brain. Cephalalgia (2019) 39(3):342–53. Epub 2017/09/01. doi: 10.1177/0333102417728873. PubMed PMID: 28856910.

66. Eftekhari S, Salvatore CA, Gaspar RC, Roberts R, O’Malley S, Zeng ZZ, et al. Localization of CGRP Receptor Components, CGRP, and Receptor Binding Sites in Human and Rhesus Cerebellar Cortex. Cerebellum (2013) 12(6):937–49. doi: 10.1007/s12311-013-0509-4. PubMed PMID: WOS:000325827300020.

67. Kawai Y, Takami K, Shiosaka S, Emson PC, Hillyard CJ, Girgis S, et al. Topographic localization of calcitonin gene-related peptide in the rat brain: an immunohistochemical analysis. Neuroscience (1985) 15(3):747–63. Epub 1985/07/01. doi: 10.1016/0306-4522(85)90076-4. PubMed PMID: 3877882.

68. Zhang XY, Wang JJ, Zhu JN. Cerebellar fastigial nucleus: from anatomic construction to physiological functions. Cerebellum Ataxias (2016) 3:9. Epub 2016/05/05. doi: 10.1186/s40673-016-0047-1. PubMed PMID: 27144010; PubMed Central PMCID: PMCPMC4853849.

69. Fujita H, Kodama T, du Lac S. Modular output circuits of the fastigial nucleus for diverse motor and nonmotor functions of the cerebellar vermis. Elife (2020) 9. Epub 2020/07/09. doi: 10.7554/eLife.58613. PubMed PMID: 32639229; PubMed Central PMCID: PMCPMC7438114.

70. Saab CY, Willis WD. Cerebellar stimulation modulates the intensity of a visceral nociceptive reflex in the rat. Exp Brain Res (2002) 146(1):117–21. Epub 2002/08/23. doi: 10.1007/s00221-002-1107-8. PubMed PMID: 12192585.

71. Zhen LL, Miao B, Chen YY, Su Z, Xu MQ, Fei SJ, et al. Protective effect and mechanism of injection of glutamate into cerebellum fastigial nucleus on chronic visceral hypersensitivity in rats. Life Sci (2018) 203:184–92. doi: 10.1016/j.lfs.2018.04.043. PubMed PMID: WOS:000432846600022.

72. Saab CY, Garcia-Nicas E, Willis WD. Stimulation in the rat fastigial nucleus enhances the responses of neurons in the dorsal column nuclei to innocuous stimuli. Neurosci Lett (2002) 327(1):17–20. Epub 2002/07/06. doi: 10.1016/s0304-3940(02)00379-8. PubMed PMID: 12098490.

73. Wattiez AS, Gaul OJ, Kuburas A, Zorilla E, Waite JS, Mason BN, et al. CGRP induces migraine-like symptoms in mice during both the active and inactive phases. J Headache Pain (2021) 22(1). doi: ARTN 62 10.1186/s10194-021-01277-9. PubMed PMID: WOS:000668540300001.

74. Mason BN, Wattiez AS, Balcziak LK, Kuburas A, Kutschke WJ, Russo AF. Vascular actions of peripheral CGRP in migraine-like photophobia in mice. Cephalalgia (2020) 40(14):1585–604. Epub 2020/08/20. doi: 10.1177/0333102420949173. PubMed PMID: 32811179; PubMed Central PMCID: PMCPMC7785273.

75. Kuburas A, Mason BN, Hing B, Wattiez AS, Reis AS, Sowers LP, et al. PACAP Induces Light Aversion in Mice by an Inheritable Mechanism Independent of CGRP. J Neurosci (2021) 41(21):4697–715. Epub 2021/04/14. doi: 10.1523/JNEUROSCI.2200-20.2021. PubMed PMID: 33846231; PubMed Central PMCID: PMCPMC8260237.

76. Wang M, Mason BN, Sowers LP, Kuburas A, Rea BJ, Russo AF. Investigating Migraine-Like Behavior using Light Aversion in Mice. JoVE (2021) (174):e62839. doi: doi:10.3791/62839.

77. Raithel SJ, Sapio MR, Iadarola MJ, Mannes AJ. Thermal A-Nociceptors, Identified by Transcriptomics, Express Higher Levels of Anesthesia-Sensitive Receptors Than Thermal C-Fibers and Are More Suppressible by Low-Dose Isoflurane. Anesth Analg (2018) 127(1):263–6. doi: 10.1213/Ane.0000000000002505. PubMed PMID: WOS:000435465900040.

78. Wattiez AS, Castonguay WC, Gaul OJ, Waite JS, Schmidt CM, Reis AS, et al. Different forms of traumatic brain injuries cause different tactile hypersensitivity profiles. Pain (2021) 162(4):1163–75. Epub 2020/10/08. doi: 10.1097/j.pain.0000000000002103. PubMed PMID: 33027220; PubMed Central PMCID: PMCPMC8008742.

79. Chaplan SR, Bach FW, Pogrel JW, Chung JM, Yaksh TL. Quantitative assessment of tactile allodynia in the rat paw. J Neurosci Methods (1994) 53(1):55–63. Epub 1994/07/01. doi: 10.1016/0165-0270(94)90144-9. PubMed PMID: 7990513.

80. Dixon WJ. The up-and-down Method for Small Samples. J Am Stat Assoc (1965) 60(312):967–78. doi: DOI 10.2307/2283398. PubMed PMID: WOS:A1965CKX1600002.

81. Rea BJ, Davison A, Ketcha M-J, Smith KJ, Fairbanks AM, Wattiez A-S, et al. Automated detection of squint as a sensitive assay of sex-dependent CGRP and amylin-induced pain in mice. PAIN (2021). doi: 10.1097/j.pain.0000000000002537. PubMed PMID: 00006396-900000000-97845.

82. Lipton RB, Bigal ME, Ashina S, Burstein R, Silberstein S, Reed ML, et al. Cutaneous allodynia in the migraine population. Ann Neurol (2008) 63(2):148–58. Epub 2007/12/07. doi: 10.1002/ana.21211. PubMed PMID: 18059010; PubMed Central PMCID: PMCPMC2729495.

83. Bigal ME, Ashina S, Burstein R, Reed ML, Buse D, Serrano D, et al. Prevalence and characteristics of allodynia in headache sufferers: a population study. Neurology (2008) 70(17):1525–33. Epub 2008/04/23. doi: 10.1212/01.wnl.0000310645.31020.b1. PubMed PMID: 18427069.

84. Langford DJ, Bailey AL, Chanda ML, Clarke SE, Drummond TE, Echols S, et al. Coding of facial expressions of pain in the laboratory mouse. Nat Methods (2010) 7(6):447–9. Epub 2010/05/11. doi: 10.1038/nmeth.1455. PubMed PMID: 20453868.

85. Rea BJ, Wattiez AS, Waite JS, Castonguay WC, Schmidt CM, Fairbanks AM, et al. Peripherally administered calcitonin gene-related peptide induces spontaneous pain in mice: implications for migraine. Pain (2018) 159(11):2306–17. Epub 2018/07/12. doi: 10.1097/j.pain.0000000000001337. PubMed PMID: 29994995; PubMed Central PMCID: PMCPMC6193822.

86. Simms J, Uddin R, Sakmar TP, Gingell JJ, Garelja ML, Hay DL, et al. Photoaffinity Cross-Linking and Unnatural Amino Acid Mutagenesis Reveal Insights into Calcitonin Gene-Related Peptide Binding to the Calcitonin Receptor-like Receptor/Receptor Activity-Modifying Protein 1 (CLR/RAMP1) Complex. Biochemistry-Us (2018) 57(32):4915–22. doi: 10.1021/acs.biochem.8b00502. PubMed PMID: WOS:000442184600015.

87. Abad N, Rosenberg JT, Hike DC, Harrington MG, Grant SC. Dynamic sodium imaging at ultra-high field reveals progression in a preclinical migraine model. Pain (2018) 159(10):2058–65. doi: 10.1097/j.pain.0000000000001307. PubMed PMID: WOS:000451227100017.

88. Jia Z, Yu S, Tang W, Zhao D. Altered functional connectivity of the insula in a rat model of recurrent headache. Mol Pain (2020) 16:1744806920922115. Epub 2020/04/28. doi: 10.1177/1744806920922115. PubMed PMID: 32338132; PubMed Central PMCID: PMCPMC7227144.

89. Jia ZH, Chen XY, Tang WJ, Zhao DF, Yu SY. Atypical functional connectivity between the anterior cingulate cortex and other brain regions in a rat model of recurrent headache. Mol Pain (2019) 15. doi: Artn 1744806919842483 10.1177/1744806919842483. PubMed PMID: WOS:000465904000001.

90. Li P, Gu H, Xu J, Zhang Z, Li F, Feng M, et al. Purkinje cells of vestibulocerebellum play an important role in acute vestibular migraine. J Integr Neurosci (2019) 18(4):409–14. Epub 2020/01/09. doi: 10.31083/j.jin.2019.04.1168. PubMed PMID: 31912699.

91. Pietrobon D. Function and dysfunction of synaptic calcium channels: insights from mouse models. Curr Opin Neurobiol (2005) 15(3):257–65. Epub 2005/06/01. doi: 10.1016/j.conb.2005.05.010. PubMed PMID: 15922581.

92. Adams PJ, Rungta RL, Garcia E, van den Maagdenberg AM, MacVicar BA, Snutch TP. Contribution of calcium-dependent facilitation to synaptic plasticity revealed by migraine mutations in the P/Q-type calcium channel. Proc Natl Acad Sci U S A (2010) 107(43):18694–9. Epub 2010/10/13. doi: 10.1073/pnas.1009500107. PubMed PMID: 20937883; PubMed Central PMCID: PMCPMC2972937.

93. Dey PK, Ray AK. Anterior cerebellum as a site for morphine analgesia and post-stimulation analgesia. Indian journal of physiology and pharmacology (1982) 26(1):3–12. Epub 1982/01/01. PubMed PMID: 7106957.

94. Siegel P, Wepsic JG. Alteration of nociception by stimulation of cerebellar structures in the monkey. Physiol Behav (1974) 13(2):189–94. Epub 1974/08/01. doi: 10.1016/0031-9384(74)90033-x. PubMed PMID: 4212925.

95. Sacchetti B, Scelfo B, Strata P. The cerebellum: synaptic changes and fear conditioning. Neuroscientist (2005) 11(3):217–27. Epub 2005/05/25. doi: 10.1177/1073858405276428. PubMed PMID: 15911871.

96. Dong WQ, Wilson OB, Skolnick MH, Dafny N. Hypothalamic, dorsal raphe and external electrical stimulation modulate noxious evoked responses of habenula neurons. Neuroscience (1992) 48(4):933–40. Epub 1992/06/01. doi: 10.1016/0306-4522(92)90281-6. PubMed PMID: 1630629.

97. Liu FY, Qiao JT, Dafny N. Cerebellar stimulation modulates thalamic noxious-evoked responses. Brain Res Bull (1993) 30(5-6):529–34. Epub 1993/01/01. doi: 10.1016/0361-9230(93)90079-q. PubMed PMID: 8457903.

98. Saab CY, Kawasaki M, Al-Chaer ED, Willis WD. Cerebellar cortical stimulation increases spinal visceral nociceptive responses. J Neurophysiol (2001) 85(6):2359–63. Epub 2001/06/02. doi: 10.1152/jn.2001.85.6.2359. PubMed PMID: 11387382.

99. Hagains CE, Senapati AK, Huntington PJ, He JW, Peng YB. Inhibition of spinal cord dorsal horn neuronal activity by electrical stimulation of the cerebellar cortex. J Neurophysiol (2011) 106(5):2515–22. Epub 2011/08/13. doi: 10.1152/jn.00719.2010. PubMed PMID: 21832034.

100. Digre KB, Brennan KC. Shedding light on photophobia. J Neuroophthalmol (2012) 32(1):68–81. Epub 2012/02/15. doi: 10.1097/WNO.0b013e3182474548. PubMed PMID: 22330853; PubMed Central PMCID: PMCPMC3485070.

101. Russo AF. Calcitonin Gene-Related Peptide (CGRP): A New Target for Migraine. Annu Rev Pharmacol (2015) 55:533–52. doi: 10.1146/annurev-pharmtox-010814-124701. PubMed PMID: WOS:000348560500028.

102. Russo AF, Recober A. Unanswered Questions in Headache: So What Is Photophobia, Anyway? Headache (2013) 53(10):1677–8. doi: 10.1111/head.12231. PubMed PMID: WOS:000327253600019.

103. Seidel S, Beisteiner R, Manecke M, Aslan TS, Wober C. Psychiatric comorbidities and photophobia in patients with migraine. J Headache Pain (2017) 18(1):18. Epub 2017/02/12. doi: 10.1186/s10194-017-0718-1. PubMed PMID: 28185159; PubMed Central PMCID: PMCPMC5307401.

104. Noseda R, Copenhagen D, Burstein R. Current understanding of photophobia, visual networks and headaches. Cephalalgia (2019) 39(13):1623–34. Epub 2018/06/27. doi: 10.1177/0333102418784750. PubMed PMID: 29940781; PubMed Central PMCID: PMCPMC6461529.

105. Martersteck EM, Hirokawa KE, Evarts M, Bernard A, Duan X, Li Y, et al. Diverse Central Projection Patterns of Retinal Ganglion Cells. Cell Rep (2017) 18(8):2058–72. Epub 2017/02/24. doi: 10.1016/j.celrep.2017.01.075. PubMed PMID: 28228269; PubMed Central PMCID: PMCPMC5357325.

106. Zwimpfer TJ, Aguayo AJ, Bray GM. Synapse formation and preferential distribution in the granule cell layer by regenerating retinal ganglion cell axons guided to the cerebellum of adult hamsters. J Neurosci (1992) 12(4):1144–59. Epub 1992/04/01. PubMed PMID: 1556590; PubMed Central PMCID: PMCPMC6575799.

107. McLean CP, Asnaani A, Litz BT, Hofmann SG. Gender differences in anxiety disorders: prevalence, course of illness, comorbidity and burden of illness. J Psychiatr Res (2011) 45(8):1027–35. Epub 2011/03/29. doi: 10.1016/j.jpsychires.2011.03.006. PubMed PMID: 21439576; PubMed Central PMCID: PMCPMC3135672.

108. Babaev O, Piletti Chatain C, Krueger-Burg D. Inhibition in the amygdala anxiety circuitry. Exp Mol Med (2018) 50(4):1–16. Epub 2018/04/10. doi: 10.1038/s12276-018-0063-8. PubMed PMID: 29628509; PubMed Central PMCID: PMCPMC5938054.

109. Teune TM, van der Burg J, van der Moer J, Voogd J, Ruigrok TJH. Topography of cerebellar nuclear projections to the brain stem in the rat. Cerebellar Modules: Molecules, Morphology, and Function (2000) 124:141–72. PubMed PMID: WOS:000180918700011.

110. Graeff FG. Serotonin, the periaqueductal gray and panic. Neurosci Biobehav Rev (2004) 28(3):239–59. Epub 2004/07/01. doi: 10.1016/j.neubiorev.2003.12.004. PubMed PMID: 15225969.

111. Maleki N, Linnman C, Brawn J, Burstein R, Becerra L, Borsook D. Her versus his migraine: multiple sex differences in brain function and structure. Brain (2012) 135(Pt 8):2546–59. Epub 2012/07/31. doi: 10.1093/brain/aws175. PubMed PMID: 22843414.

112. Benatto MT, Florencio LL, Carvalho GF, Dach F, Bigal ME, Chaves TC, et al. Cutaneous allodynia is more frequent in chronic migraine, and its presence and severity seems to be more associated with the duration of the disease. Arq Neuro-Psiquiat (2017) 75(3):153–9. doi: 10.1590/0004-282x20170015. PubMed PMID: WOS:000397873200005.

113. Louter MA, Bosker JE, van Oosterhout WPJ, van Zwet EW, Zitman FG, Ferrari MD, et al. Cutaneous allodynia as a predictor of migraine chronification. Brain (2013) 136:3489–96. doi: 10.1093/brain/awt251. PubMed PMID: WOS:000326291800029.

114. Burstein R, Yarnitsky D, Goor-Aryeh I, Ransil BJ, Bajwa ZH. An association between migraine and cutaneous allodynia. Annals of Neurology (2000) 47(5):614–24. doi: DOI 10.1002/1531-8249(200005)47:5<614::Aid-Ana9>3.3.Co;2-E. PubMed PMID: WOS:000086731000009.

115. Avona A, Burgos-Vega C, Burton MD, Akopian AN, Price TJ, Dussor G. Dural Calcitonin Gene-Related Peptide Produces Female-Specific Responses in Rodent Migraine Models. J Neurosci (2019) 39(22):4323–31. Epub 2019/04/10. doi: 10.1523/JNEUROSCI.0364-19.2019. PubMed PMID: 30962278; PubMed Central PMCID: PMCPMC6538861.

116. Kao CH, Wang SJ, Tsai CF, Chen SP, Wang YF, Fuh JL. Psychiatric comorbidities in allodynic migraineurs. Cephalalgia (2014) 34(3):211–8. Epub 2013/09/21. doi: 10.1177/0333102413505238. PubMed PMID: 24048892.

117. Han SM, Kim KM, Cho SJ, Yang KI, Kim D, Yun CH, et al. Prevalence and characteristics of cutaneous allodynia in probable migraine. Sci Rep (2021) 11(1):2467. Epub 2021/01/30. doi: 10.1038/s41598-021-82080-z. PubMed PMID: 33510340; PubMed Central PMCID: PMCPMC7844001.

118. Wang M, Wattiez A-S, Russo AF. CGRP Antibodies for Animal Models of Primary and Secondary Headache Disorders. In: Maassen van den Brink A, Martelletti P, editors. Monoclonal Antibodies in Headache : From Bench to Patient. Cham: Springer International Publishing (2021). p. 69–97.

119. Ruscheweyh R, Kuhnel M, Filippopulos F, Blum B, Eggert T, Straube A. Altered experimental pain perception after cerebellar infarction. Pain (2014) 155(7):1303–12. doi: 10.1016/j.pain.2014.04.006. PubMed PMID: WOS:000339136100020.

120. van den Pol AN. Neuropeptide transmission in brain circuits. Neuron (2012) 76(1):98–115. Epub 2012/10/09. doi: 10.1016/j.neuron.2012.09.014. PubMed PMID: 23040809; PubMed Central PMCID: PMCPMC3918222.

121. Hay DL, Walker CS. CGRP and its receptors. Headache (2017) 57(4):625–36. Epub 2017/02/25. doi: 10.1111/head.13064. PubMed PMID: 28233915.

