## Supplemental Figures for "CGRP Administration into the Cerebellum Evokes Migraine-like Behaviors Predominately in Female Mice"

Supplementary Material

### 1.1 Supplementary Figures

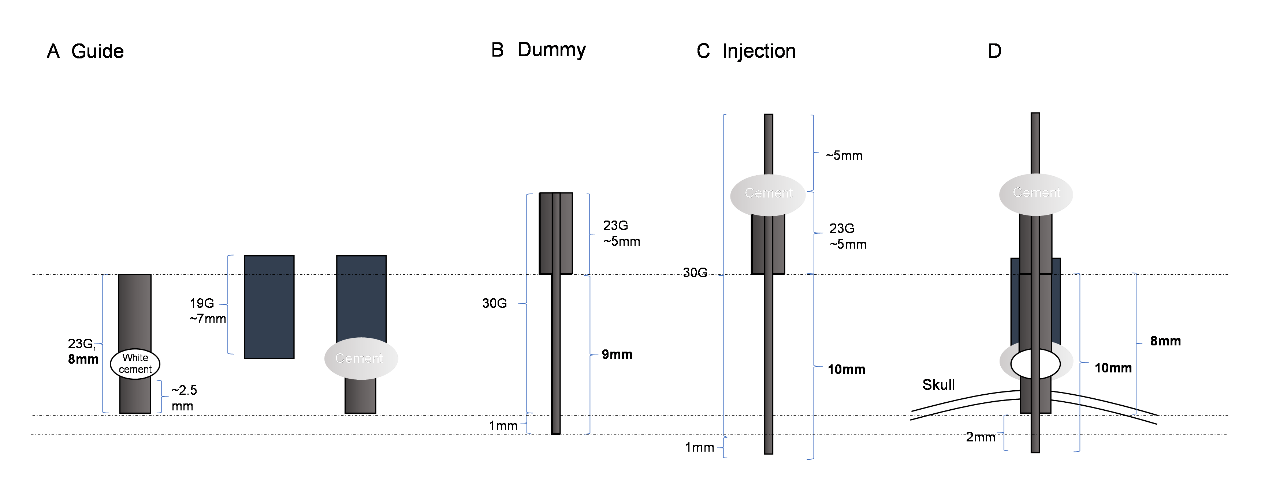

**Supplementary Fig. 1 Cannula design and assembly diagram.** (**A**) Guide cannulas. The guide cannula was made from an 8-mm, 23-gauge needle (left) with the ventral portion covered by a ~7-mm, 19-gauge tubing (middle), which were adhered by adhesive and dental cement (right). The plastic holder was removed from 23-gauge needle without removing the original white cement (left). The ~7-mm, 19-gauge tubing is ~2 mm higher than the 23-guage needle to shield the junction between guide’s top and the dummy or injection cannula after their insertion, allowing the dummy or the injection cannula to stay in position after insertion (right). (**B**) Dummy cannulas. The dummy cannula, used to seal and keep the guide cannula free of clogs, was made by crimping a short segment of ~5-mm, 23-gauge tubing to a ~14 mm piece of 30-gauge tubing. The bottom of the 30-gauge tubing was cut to ensure that the 30-gauge segment below the ~5-mm, 23-gauge segment is 9 mm. (**C**) Injection cannulas. The injection cannula was made by adhering a short segment of ~5-mm, 23-gauge tubing ~5 mm below the top of a ~20-mm piece of 30-gauge tubing with adhesive and dental cement. The bottom of the 30-gauge tubing was cut to ensure that the 30-gauge segment below the ~5-mm, 23-gauge segment is 10 mm. (**D**) The injection cannula extended 2 mm beyond the base of the guide cannula when was inserted into the guide cannula.

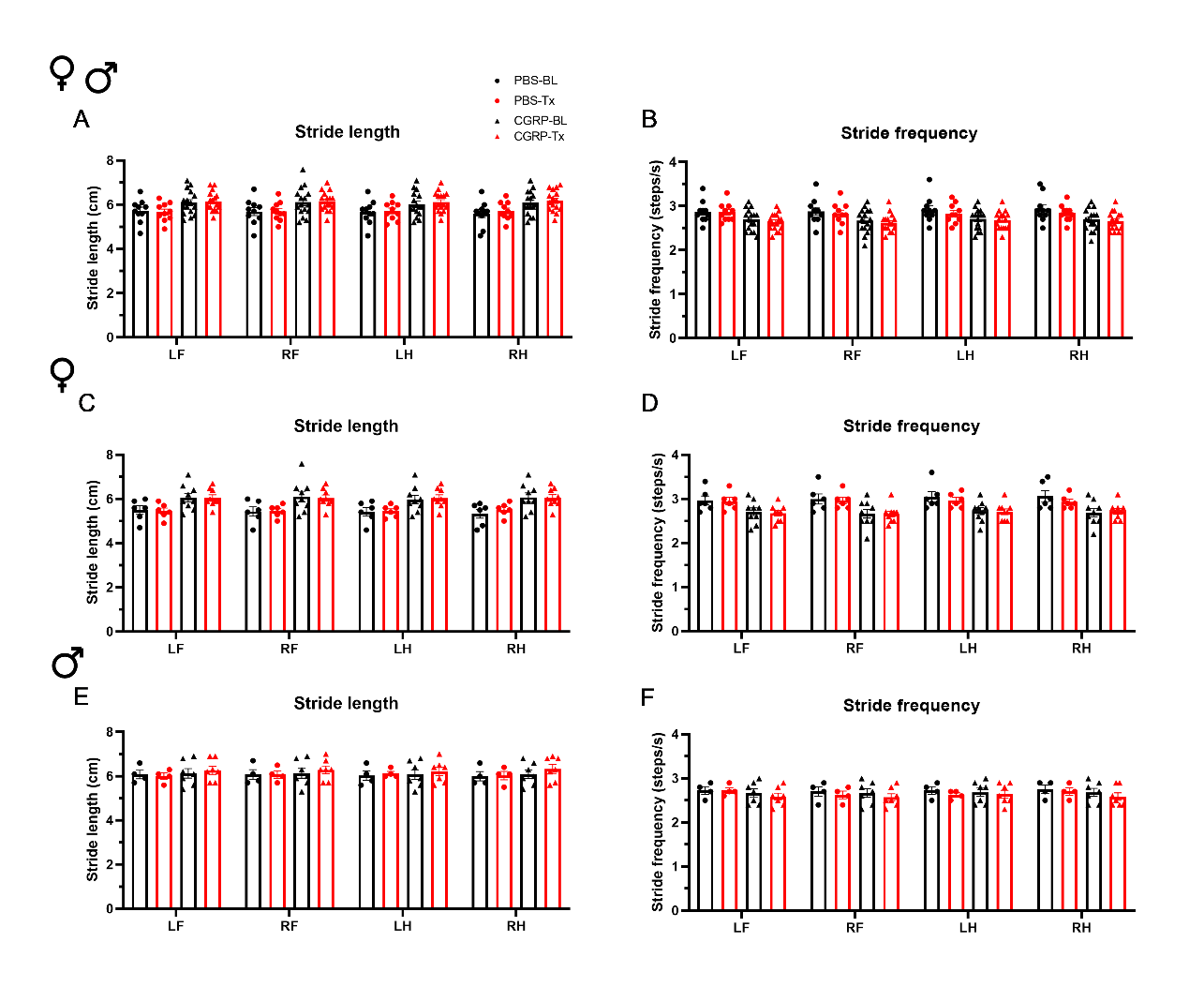

**Supplementary Fig. 2 CGRP injection into the MN did not induce gait alterations.** Stride length **(A)** and frequency **(B)** for all mice following injection of PBS (n=10) or CGRP (1 μg/200 nl; n=16) into the right MN of C57BL/6J mice via cannulas. Stride length **(C)** and frequency **(D)** for female mice (PBS: n=6; CGRP: n=9). Stride length **(E)** and frequency **(F)** for male mice (PBS: n=4; CGRP: n=7). Injection of CGRP into the MN did not change the stride length and frequency comparing before and after CGRP or PBS treatments across and within sexes. It suggests that CGRP in the MN did not change the gait. LF: left front paw; RF: right front paw; LH: left hind paw; RH: right hind paw. Data are the mean ± SEM. Statistics are described in Supplementary Table 1. Data are from one experiment.

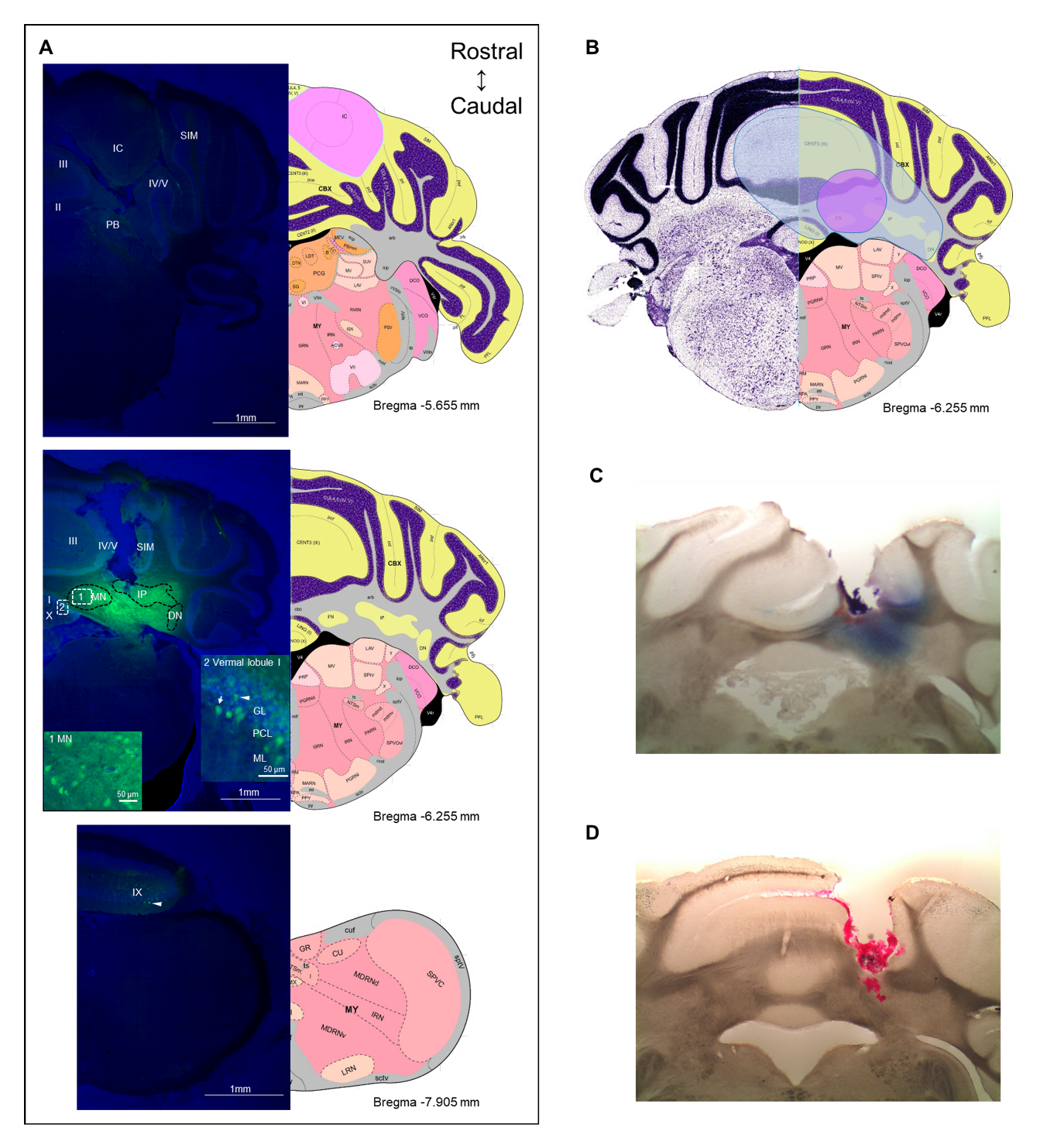

**Supplementary Fig. 3 The diffusion range of CGRP. (A)** Representative example of a mouse after injection of fluorescein-15-CGRP. **Upper panel:** In the most rostral section, dim fluorescence was detected in the inferior colliculus, the parabrachial nucleus, vermal lobules II-V, and the simple lobule. **Middle panel:** Fluorescein-15-CGRP at the injection site. Areas within rectangle are magnified in boxes 1 and 2. Clusters of fluorescein-15-CGRP were detected in cell bodies in the MN (box 1) and nearby cells, including the interposed and lateral cerebellar nuclei, granular, Purkinje cell, and molecular layers of vermal lobules I/III/IV/V (box 2). Dim signal was found in the simple lobule of the hemispheric regions. **Lower panel:** In the most caudal section, dim fluorescence was detected in the lobule IX. Green: fluorescein-15-CGRP; Blue: TO-PRO-3**. (B)** The spread of the green fluorescence among the mice injected with fluorescein-15-CGRP. The smallest (purple shading) spread of signals covers the MN and few of nearby cells in the vermal lobules III/IV/V. The largest spread (blue shading) covers the MN and cells beyond the MN including vermal lobules I/III/IV/V/X, the simple lobule and other cerebellar deep nuclei. In summary, florescent signals were found in vermal lobules I-X, the simple lobule in the hemispheric region, and the midbrain (mainly in superior and inferior colliculus) from rostrally to caudally. (**C**) A representative image of a mouse with injection of Evans blue. (**D**) A representative image of a mouse with injection of red beads. DN: lateral cerebellar nucleus (dentate nucleus); GL: granular layer; IC: inferior colliculus; IP: interposed nucleus; ML: molecular layer; MN: medial cerebellar nucleus; PB: parabrachial nucleus; PCL: Purkinje cell layer; SIM: simple lobule. Image credit: Allen Institute. Numbers indicate the distance from bregma in the anteroposterior plane in Allen Mouse Brain Atlas coronal images.

**Supplementary Table 1 Statistical analyses**

| Figure no. | Analysis | Statistics (symbol on Figure) | | | |
| --- | --- | --- | --- | --- | --- |
| Fig. 1A left (all mice) | Two-way repeated measure ANOVA | | | | |
|  | Interaction factor | F (5, 95) = 1.734, P=0.1343 | | | |
|  | Treatment factor | F (1, 19) = 23.23, P=0.0001 | | | |
|  | Time factor | F (2.677, 50.86) = 6.280, P=0.0015 | | | |
| Fig. 1A middle (females) | Two-way repeated measure ANOVA | | | | |
|  | Interaction factor | F (5, 40) = 0.7746, P=0.5738 | | | |
|  | Treatment factor | F (1, 8) = 14.45, P=0.0052 | | | |
|  | Time factor | F (2.675, 21.40) = 9.368, P=0.0005 | | | |
| Fig. 1A right (males) | Two-way repeated measure ANOVA | | | | |
|  | Interaction factor | F (5, 45) = 1.074, P=0.3877 | | | |
|  | Treatment factor | F (1, 9) = 8.808, P=0.0158 | | | |
|  | Time factor | F (2.076, 18.68) = 1.891, P=0.1777 | | | |
| Fig. 1B left (all mice) | Unpaired 2-tailed t-test | t=4.820, df=19, P=0.0001 | | | |
| Fig. 1B middle (females) | Unpaired 2-tailed t-test | t=3.802, df=8, P=0.0052 | | | |
| Fig. 1B right (males) | Unpaired 2-tailed t-test | t=2.968, df=9, P=0.0158 | | | |
| Fig. 2A upper panel |  |  | | | |
| Left (all mice) | Two-way repeated measure ANOVA (mixed effects analysis) | | | | |
|  | Interaction factor | F (5, 84) = 0.7322, P=0.6013 | | | |
|  | Treatment factor | F (1, 19) = 0.01297, P=0.9105 | | | |
|  | Time factor | F (2.807, 47.16) = 8.859, P=0.0001 | | | |
| Middle (females) | Two-way repeated measure ANOVA (mixed effects analysis) | | | | |
|  | Interaction factor | F (5, 36) = 0.4571, P=0.8053 | | | |
|  | Treatment factor | F (1, 8) = 0.6466, P=0.4446 | | | |
|  | Time factor | F (1.952, 14.05) = 7.334, P=0.0068 | | | |
| Right (males) | Two-way repeated measure ANOVA (mixed effects analysis) | | | | |
|  | Interaction factor | F (5, 38) = 1.570, P=0.1918 | | | |
|  | Treatment factor | F (1, 9) = 0.7938, P=0.3961 | | | |
|  | Time factor | F (1.714, 13.03) = 6.755, P=0.0119 | | | |
| Fig. 2A lower panel |  |  | | | |
| Left (all mice) | Two-way repeated measure ANOVA | | | | |
|  | Interaction factor | F (5, 95) = 2.995, P=0.0148 | | | |
|  | Treatment factor | F (1, 19) = 28.00, P<0.0001 | | | |
|  | Time factor | F (3.220, 61.18) = 41.21, P<0.0001 | | | |
|  | Šídák's multiple comparisons test | *P < .05, **P < .01, ***P < .001 | | | |
| Middle (females) | Two-way repeated measure ANOVA | | | | |
|  | Interaction factor | F (5, 40) = 4.713, P=0.0018 | | | |
|  | Treatment factor | F (1, 8) = 17.02, P=0.0033 | | | |
|  | Time factor | F (2.779, 22.23) = 31.66, P<0.0001 | | | |
|  | Šídák's multiple comparisons test | *P < .05, **P < .01 | | | |
| Right (males) | Two-way repeated measure ANOVA | | | | |
|  | Interaction factor | F (5, 45) = 0.6068, P=0.6950 | | | |
|  | Treatment factor | F (1, 9) = 10.24, P=0.0108 | | | |
|  | Time factor | F (2.691, 24.22) = 15.43, P<0.0001 | | | |
| Fig. 2B upper panel |  |  | | | |
| Left (all mice) | Unpaired 2-tailed t-test | t=0.06504, df=19, P=0.9488 | | | |
| Middle (females) | Unpaired 2-tailed t-test | t=0.9003, df=8, P=0.3942 | | | |
| Right (males) | Unpaired 2-tailed t-test | t=0.7370, df=9, P=0.4799 | | | |
| Fig. 2B lower panel | Unpaired 2-tailed t-test |  | | | |
| Left (all mice) | Unpaired 2-tailed t-test | t=5.291, df=19, P<0.0001 | | | |
| Middle (females) | Unpaired 2-tailed t-test | t=4.125, df=8, P=0.0033 | | | |
| Right (males) | Unpaired 2-tailed t-test | t=3.200, df=9, P=0.0108 | | | |
| Fig. 2C upper panel |  |  | | | |
| Left (all mice) | Two-way repeated measure ANOVA (mixed effects analysis) | | | | |
|  | Interaction factor | F (5, 84) = 1.411, P=0.2288 | | | |
|  | Treatment factor | F (1, 19) = 4.694, P=0.0432 | | | |
|  | Time factor | F (2.759, 46.35) = 0.5223, P=0.6542 | | | |
| Middle (females) | Two-way repeated measure ANOVA (mixed effects analysis) | | | | |
|  | Interaction factor | F (5, 36) = 2.016, P=0.0997 | | | |
|  | Treatment factor | F (1, 8) = 3.530, P=0.0971 | | | |
|  | Time factor | F (1.482, 10.67) = 0.1453, P=0.8048 | | | |
| Right (males) | Two-way repeated measure ANOVA (mixed effects analysis) | | | | |
|  | Interaction factor | F (5, 38) = 0.9688, P=0.4491 | | | |
|  | Treatment factor | F (1, 9) = 1.575, P=0.2411 | | | |
|  | Time factor | F (1.905, 14.48) = 0.9849, P=0.3933 | | | |
| Fig. 2C lower panel |  |  | | | |
| Left (all mice) | Two-way repeated measure ANOVA | | | | |
|  | Interaction factor | F (5, 95) = 1.020, P=0.4104 | | | |
|  | Treatment factor | F (1, 19) = 4.554, P=0.0461 | | | |
|  | Time factor | F (1.690, 32.10) = 5.394, P=0.0129 | | | |
| Middle (females) | Two-way repeated measure ANOVA | | | | |
|  | Interaction factor | F (5, 40) = 2.197, P=0.0737 | | | |
|  | Treatment factor | F (1, 8) = 3.328, P=0.1055 | | | |
|  | Time factor | F (1.459, 11.67) = 4.477, P=0.0453 | | | |
| Right (males) | Two-way repeated measure ANOVA | | | | |
|  | Interaction factor | F (5, 45) = 0.4386, P=0.8192 | | | |
|  | Treatment factor | F (1, 9) = 1.189, P=0.3039 | | | |
|  | Time factor | F (2.333, 20.99) = 2.168, P=0.1332 | | | |
| Fig. 2D upper panel |  |  | | | |
| Left (all mice) | Unpaired 2-tailed t-test | t=2.300, df=19, P=0.0330 | | | |
| Middle (females) | Unpaired 2-tailed t-test | t=1.979, df=8, P=0.0831 | | | |
| Right (males) | Unpaired 2-tailed t-test | t=1.321, df=9, P=0.2192 | | | |
| Fig. 2D lower panel | Unpaired 2-tailed t-test |  | | | |
| Left (all mice) | Unpaired 2-tailed t-test | t=2.131, df=19, P=0.0464 | | | |
| Middle (females) | Unpaired 2-tailed t-test | t=1.854, df=8, P=0.1008 | | | |
| Right (males) | Unpaired 2-tailed t-test | t=1.084, df=9, P=0.3066 | | | |
| Fig. 2E |  |  | | | |
| Left (all mice) | Two-way repeated measure ANOVA | | | | |
|  | Interaction factor | F (5, 95) = 2.190, P=0.0617 | | | |
|  | Treatment factor | F (1, 19) = 19.19, P=0.0003 | | | |
|  | Time factor | F (3.507, 66.63) = 7.639, P<0.0001 | | | |
| Middle (females) | Two-way repeated measure ANOVA | | | | |
|  | Interaction factor | F (5, 40) = 2.181, P=0.0755 | | | |
|  | Treatment factor | F (1, 8) = 10.05, P=0.0132 | | | |
|  | Time factor | F (2.947, 23.58) = 4.349, P=0.0145 | | | |
| Right (males) | Two-way repeated measure ANOVA | | | | |
|  | Interaction factor | F (5, 45) = 0.6413, P=0.6694 | | | |
|  | Treatment factor | F (1, 9) = 8.275, P=0.0183 | | | |
|  | Time factor | F (2.478, 22.31) = 3.553, P=0.0375 | | | |
| Fig. 2F |  |  | | | |
| Left (all mice) | Unpaired 2-tailed t-test | t=4.380, df=19, P=0.0003 | | | |
| Middle (females) | Unpaired 2-tailed t-test | t=3.170, df=8, P=0.0132 | | | |
| Right (males) | Unpaired 2-tailed t-test | t=2.877, df=9, P=0.0183 | | | |
| Fig. 3A left (all mice) | Two-way repeated measure ANOVA | | | | |
|  | Interaction factor | F (5, 100) = 5.414, P=0.0002 | | | |
|  | Treatment factor | F (1, 20) = 9.155, P=0.0067 | | | |
|  | Time factor | F (3.129, 62.59) = 9.223, P<0.0001 | | | |
|  | Šídák's multiple comparisons test | *P < .05, **P < .01 | | | |
| Fig. 3A middle (females) | Two-way repeated measure ANOVA | | | | |
|  | Interaction factor | F (5, 40) = 1.270, P=0.2960 | | | |
|  | Treatment factor | F (1, 8) = 8.899, P=0.0175 | | | |
|  | Time factor | F (3.397, 27.17) = 3.397, P=0.0276 | | | |
| Fig. 3A right (males) | Two-way repeated measure ANOVA | | | | |
|  | Interaction factor | F (5, 50) = 4.846, P=0.0011 | | | |
|  | Treatment factor | F (1, 10) = 4.013, P=0.0730 | | | |
|  | Time factor | F (2.374, 23.74) = 5.795, P=0.0065 | | | |
| Fig. 3B left (all mice) | Unpaired 2-tailed t-test | t=3.026, df=20, P=0.0067 | | | |
| Fig. 3B middle (females) | Unpaired 2-tailed t-test | t=2.983, df=8, P=0.0175 | | | |
| Fig. 3B right (males) | Unpaired 2-tailed t-test | t=2.003, df=10, P=0.0730 | | | |
| Fig. 4A left (all mice) | Two-way repeated measure ANOVA | | | | |
|  | Interaction factor | F (1, 41) = 3.684, P=0.0619 | | | |
|  | Treatment factor | F (1, 41) = 0.003699, P=0.9518 | | | |
|  | Condition factor | F (1, 41) = 17.87, P=0.0001 | | | |
|  | Šídák's multiple comparisons test | ****P < 0.0001 | | | |
| Fig. 4A middle (females) | Two-way repeated measure ANOVA | | | | |
|  | Interaction factor | F (1, 20) = 8.429, P=0.0088 | | | |
|  | Treatment factor | F (1, 20) = 0.06922, P=0.7952 | | | |
|  | Condition factor | F (1, 20) = 9.630, P=0.0056 | | | |
|  | Šídák's multiple comparisons test | ***P < .001 | | | |
| Fig. 4A right (males) | Two-way repeated measure ANOVA | | | | |
|  | Interaction factor | F (1, 19) = 0.1018, P=0.7532 | | | |
|  | Treatment factor | F (1, 19) = 0.05351, P=0.8195 | | | |
|  | Condition factor | F (1, 19) = 8.113, P=0.0103 | | | |
| Fig. 4B left (all mice) | Two-way repeated measure ANOVA | | | | |
|  | Interaction factor | F (1, 41) = 0.01288, P=0.9102 | | | |
|  | Treatment factor | F (1, 41) = 0.005324, P=0.9422 | | | |
|  | Condition factor | F (1, 41) = 44.42, P<0.0001 | | | |
|  | Šídák's multiple comparisons test | ***P < .001, ****P < .0001 | | | |
| Fig. 4B middle (females) | Two-way repeated measure ANOVA | | | | |
|  | Interaction factor | F (1, 20) = 0.1767, P=0.6787 | | | |
|  | Treatment factor | F (1, 20) = 0.7693, P=0.3908 | | | |
|  | Condition factor | F (1, 20) = 14.41, P=0.0011 | | | |
|  | Šídák's multiple comparisons test | **P < .01 | | | |
| Fig. 4B right (males) | Two-way repeated measure ANOVA | | | | |
|  | Interaction factor | F (1, 19) = 0.2071, P=0.6542 | | | |
|  | Treatment factor | F (1, 19) = 1.028, P=0.3233 | | | |
|  | Condition factor | F (1, 19) = 30.49, P<0.0001 | | | |
|  | Šídák's multiple comparisons test | **P < .01 | | | |
| Fig. 5A right (all mice) | Two-way repeated measure ANOVA | | | | |
|  | Interaction factor | F (1, 55) = 4.902, P=0.0310 | | | |
|  | Treatment factor | F (1, 55) = 0.7006, P=0.4062 | | | |
|  | Condition factor | F (1, 55) = 10.26, P=0.0023 | | | |
|  | Šídák's multiple comparisons test | ***P < .001 | | | |
| Fig. 5B right (females) | Two-way repeated measure ANOVA | | | | |
|  | Interaction factor | F (1, 29) = 2.756, P=0.1077 | | | |
|  | Treatment factor | F (1, 29) = 0.08961, P=0.7668 | | | |
|  | Condition factor | F (1, 29) = 9.036, P=0.0054 | | | |
|  | Šídák's multiple comparisons test | **P < .01 | | | |
| Fig. 5C right (males) | Two-way repeated measure ANOVA | | | | |
|  | Interaction factor | F (1, 24) = 2.309, P=0.1417 | | | |
|  | Treatment factor | F (1, 24) = 0.8393, P=0.3687 | | | |
|  | Condition factor | F (1, 24) = 3.282, P=0.0826 | | | |
|  |  | LF | RF | LH | RH |
| Suppl. Fig. 2A (all mice) | Two-way repeated measure ANOVA | | | | |
|  | Interaction factor | F (1, 24) = 0.6154, P=0.4404 | F (1, 24) = 0.01717, P=0.8968 | F (1, 24) = 0.08938, P=0.7675 | F (1, 24) = 0.02838, P=0.8676 |
|  | Treatment factor | F (1, 24) = 5.359, P=0.0295 | F (1, 24) = 4.852, P=0.0375 | F (1, 24) = 4.018, P=0.0564 | F (1, 24) = 6.202, P=0.0201 |
|  | Condition factor | F (1, 24) = 0.0006404, P=0.9800 | F (1, 24) = 0.06471, P=0.8014 | F (1, 24) = 1.430, P=0.2434 | F (1, 24) = 1.882, P=0.1828 |
| Suppl. Fig. 2B (all mice) | Two-way repeated measure ANOVA | | | | |
|  | Interaction factor | F (1, 24) = 0.5979, P=0.4469 | F (1, 24) = 2.546e-029, P>0.9999 | F (1, 24) = 1.158, P=0.2926 | F (1, 24) = 1.026, P=0.3213 |
|  | Treatment factor | F (1, 26) = 5.467, P=0.0280 | F (1, 24) = 5.044, P=0.0342 | F (1, 24) = 4.075, P=0.0548 | F (1, 24) = 5.840, P=0.0236 |
|  | Condition factor | F (1, 24) = 1.227, P=0.2790 | F (1, 24) = 1.748, P=0.1986 | F (1, 24) = 3.624, P=0.0690 | F (1, 24) = 2.849, P=0.1044 |
| Suppl. Fig. 2C (females) | Two-way repeated measure ANOVA | | | | |
|  | Interaction factor | F (1, 13) = 0.04835, P=0.8294 | F (1, 13) = 0.1587, P=0.6969 | F (1, 13) = 0.04021, P=0.8442 | F (1, 13) = 1.409, P=0.2565 |
|  | Treatment factor | F (1, 13) = 6.555, P=0.0237 | F (1, 13) = 6.113, P=0.0280 | F (1, 13) = 6.307, P=0.0260 | F (1, 13) = 7.275, P=0.0183 |
|  | Condition factor | F (1, 13) = 0.1194, P=0.7352 | F (1, 13) = 0.05712, P=0.8148 | F (1, 13) = 0.3619, P=0.5578 | F (1, 13) = 0.8653, P=0.3692 |
| Suppl. Fig. 2D (females) | Two-way repeated measure ANOVA | | | | |
|  | Interaction factor | F (1, 13) = 0.04216, P=0.8405 | F (1, 13) = 0.05000, P=0.8265 | F (1, 13) = 0.7695, P=0.3963 | F (1, 13) = 3.145, P=0.0996 |
|  | Treatment factor | F (1, 13) = 5.438, P=0.0364 | F (1, 13) = 7.020, P=0.0200 | F (1, 13) = 7.021, P=0.0200 | F (1, 13) = 6.418, P=0.0250 |
|  | Condition factor | F (1, 13) = 0.3795, P=0.5485 | F (1, 13) = 0.2000, P=0.6621 | F (1, 13) = 1.316, P=0.2720 | F (1, 13) = 1.132, P=0.3067 |
| Suppl. Fig. 2E (males) | Two-way repeated measure ANOVA | | | | |
|  | Interaction factor | F (1, 9) = 0.6454, P=0.4425 | F (1, 9) = 0.3455, P=0.5711 | F (1, 9) = 0.03195, P=0.8621 | F (1, 9) = 0.5894, P=0.4623 |
|  | Treatment factor | F (1, 9) = 0.3741, P=0.5559 | F (1, 9) = 0.2286, P=0.6439 | F (1, 9) = 0.06064, P=0.8110 | F (1, 9) = 0.5216, P=0.4885 |
|  | Condition factor | F (1, 9) = 0.06262, P=0.8080 | F (1, 9) = 0.3455, P=0.5711 | F (1, 9) = 1.026, P=0.3376 | F (1, 9) = 0.8910, P=0.3698 |
| Suppl. Fig. 2F (males) | Two-way repeated measure ANOVA | | | | |
|  | Interaction factor | F (1, 9) = 0.8070, P=0.3924 | F (1, 9) = 0.04120, P=0.8437 | F (1, 9) = 0.3445, P=0.5717 | F (1, 9) = 0.1848, P=0.6774 |
|  | Treatment factor | F (1, 9) = 0.6519, P=0.4403 | F (1, 9) = 0.1070, P=0.7511 | F (1, 9) = 0.007599, P=0.9324 | F (1, 9) = 0.5434, P=0.4798 |
|  | Condition factor | F (1, 9) = 0.8070, P=0.3924 | F (1, 9) = 2.019, P=0.1891 | F (1, 9) = 2.153, P=0.1763 | F (1, 9) = 1.663, P=0.2294 |
